# Region-dependent regulation of Tau phosphorylation by Glycerophosphodiester Phosphodiesterase 2 in a mouse model of tauopathy

**DOI:** 10.64898/2026.09.14.751492

**Authors:** Consuelo Jimenez-Ornelas, Shanthini Sockanathan

**Affiliations:** The Solomon Snyder Department of Neuroscience, The Johns Hopkins School of Medicine, Baltimore, Maryland, United States of America

## Abstract

Hyperphosphorylation of Tau promotes its aggregation and neurofibrillary tangle (NFT) formation, contributing to neuronal dysfunction and neurodegeneration in diseases such as Alzheimer’s Disease (AD) and AD-related dementias (ADRDs). However, the mechanisms underlying dysregulated Tau phosphorylation under pathological contexts remain unclear. Glycerophosphodiester phosphodiesterase 2 (GDE2) is a six-transmembrane enzyme that acts at the cell surface to cleave the glycosylphosphatidylinositol (GPI)-anchor that tethers a subclass of proteins to the membrane. Here, we show that in the *PS19* tauopathy mouse model, GDE2 disruption modulates Tau phosphorylation by transiently decreasing Tau’s propensity for aggregation through region-specific regulation of local kinase environments. In the cortex, GDE2 ablation in 6-month-old *PS19* male mice (*PS19;Gde2KO*) reduces Tau phosphorylation at Serine (S)202/Threonine (T)205 and T212 but increases S262 phosphorylation in the hippocampus of 6-month-old *PS19;Gde2KO* male and female mice. Consistent with these changes, in males, Glycogen Synthase Kinase-3α/β (GSK3α/β) activity is decreased in the cortex, while AKT activity is increased in both males and females in the hippocampus, with no changes in protein phosphatase 1 (PP1) or protein phosphatase 2A (PP2A) activity. Primary cortical neurons from *PS19;Gde2KO* animals showed reduced Tau phosphorylation at S202/T205, implying cell-autonomous roles for neuronal GDE2 in this process. GDE2 overexpression in heterologous SH-SY5Y cells increased Tau phosphorylation at S202/T205, while a catalytically inactive form of GDE2 did not, suggesting that GDE2 regulation of target GPI-anchored protein surface activity is required to modulate Tau phosphorylation. Taken together, our study identifies GDE2 as a component of the complex regulatory network that controls Tau phosphorylation in the context of tauopathy and provides insight into putative pathways relevant to Tau pathologies observed in disease.

## Introduction

Tau is a microtubule-associated protein that is encoded by the *MAPT* gene and primarily expressed in neurons, where it mainly localizes to axons [1,2]. There, Tau directly binds to microtubules, promotes their polymerization, and limits their dissociation, stabilizing the microtubule lattice essential for axonal transport [3–7]. Tau function is dynamically regulated by several post-translational modifications (PTMs), of which phosphorylation is one of the most well-studied and prominent PTMs modulating Tau function [8–10]. In disease, Tau phosphorylation homeostasis is dysregulated, leading to Tau hyperphosphorylation, aggregation, and dysfunction that ultimately contribute to neurodegeneration [10–13]. Despite the established links between dysregulated Tau phosphorylation and neurodegeneration, the specific mechanisms that regulate site-specific and temporal patterns of Tau phosphorylation in disease are not fully understood [10,14]. Identifying these regulatory mechanisms is important for furthering our understanding of the molecular basis of tauopathy progression and for developing therapeutic strategies that intervene before Tau pathology becomes established.

Tau phosphorylation is a highly dynamic process maintained by a balance between kinases and phosphatases that, respectively, phosphorylate and dephosphorylate Tau at specific residues [15,16]. Tau can be phosphorylated by a variety of kinases, including tyrosine kinases, proline-directed serine/threonine kinases, and non-proline-directed serine/threonine kinases [15–17]. While some kinases such as GSK3α/β phosphorylate Tau directly at multiple residues, other kinases, such as AKT, can phosphorylate Tau directly at select residues and indirectly regulate Tau phosphorylation by modulating the activity of other kinases, including GSK3α/β [15,17–20]. Similarly, multiple phosphatases can dephosphorylate Tau [15,21]. Although PP2A, PP1, protein phosphatase 5 (PP5), and protein phosphatase 2B (PP2B) are all able to dephosphorylate Tau with different affinities at various residues, PP2A is the main phosphatase responsible for regulating Tau phosphorylation [21].

Under physiological conditions, Tau has approximately two to three phosphate groups per molecule [22]. However, under pathological conditions, this number can increase three-to fourfold and is believed to arise from an imbalance between kinase and phosphatase activity [11,22]. Notably, hyperphosphorylation at different Tau residues can result in differential and even competing effects on Tau’s ability to bind microtubules, aggregate, and undergo proteolytic degradation [10,23,24]. For example, phosphorylation at S262 greatly reduces Tau’s affinity for microtubules, resulting in microtubule detachment [24–26]. Accordingly, S262 phosphorylation can mitigate aggregation (anti-aggregation) but can also be considered pro-seeding in the context of disease, due to resultant increases in the overall amount of cytoplasmic Tau [24–27]. Other sites of phosphorylation known to promote Tau aggregation (pro-aggregation) include T212 and S202/T205, which moderately reduce Tau’s affinity for microtubules, and S396, which, in addition to reducing microtubule affinity, also protects Tau from proteolytic degradation [10,19,24,27–30].

Given the complex network involved in modulating Tau phosphorylation, our understanding of the mechanisms underlying Tau phosphorylation dynamics is still limited; as such, identifying novel modulators is of particular interest. GDE2 is one of three six-transmembrane proteins that act at the cell surface to cleave the GPI-anchor that tethers some proteins to the cell membrane [31], and it is the only one to do so in neurons [31,32]. GDE2 is implicated in contributing to disease pathology in AD and ADRDs such as amyotrophic lateral sclerosis/frontotemporal dementia (ALS/FTD) [33,34]. Notably, GDE2 aberrantly accumulates intracellularly in the neurons of patients with AD and patients with ALS, and there is a disproportionate reduction of released GPI-anchored proteins in the cerebrospinal fluid of patients with ALS [33,34]. Studies modeling GDE2 loss-of-function demonstrate age-progressive cognitive and motor deficits in *Gde2* knockout (KO) mice [35], as well as increased production of amyloid-beta (Aβ)42 and downregulation and mislocalization of TAR DNA-binding protein 43 (TDP-43) [34,36,37], a pathology linked to neurodegeneration across AD and ADRDs. Functional studies reveal that these cellular abnormalities result from increased surface expression and activity of select GPI-anchored proteins due to impaired GDE2 GPI-anchor cleavage function in neurons [34,37]. Interestingly, ablation of GDE2 in the *APP/PS1* mouse model of amyloidosis not only exacerbated Aβ pathology but also increased Tau phosphorylation at S202/T205 [34], thus raising the question of whether GDE2 is involved in modulating Tau hyperphosphorylation, or whether this effect is limited to an amyloid-driven context. To address this, we assessed GDE2’s role in Tau phosphorylation independent of amyloid pathology.

The *PS19* humanized tauopathy mouse model, which carries a *P301S* mutation in the human *MAPT* gene driven by the mouse prion-protein promoter, is a useful animal model for studying the pathways underlying Tau hyperphosphorylation and associated Tau pathologies [38]. Although the *P301S* mutation is causal for familial FTD, the *PS19* mouse model is used as a general tauopathy model because these animals exhibit cognitive deficits and age-progressive Tau hyperphosphorylation, NFT accumulation, and neurodegeneration, similar to those seen in tauopathies like AD [38–43]. At 1.5 months, Tau seeding activity can be detected in these animals, well before Tau pathology is observed [41]. By 3 months of age, Tau hyperphosphorylation has initiated, and gliosis and synaptic loss can be observed in the hippocampus [38]. By 6 months, gliosis and Tau hyperphosphorylation further increase and spread to the amygdala and entorhinal cortex, synaptic loss worsens and spreads to cortical regions, and NFTs begin to accumulate in the hippocampus, amygdala, and entorhinal cortex [38]. Cognitive impairments, such as impaired spatial learning and memory, are also evident [40]. Finally, by 9 months, NFT pathology becomes more severe, spreading to cortical regions, and robust neuronal loss is evident, particularly in the hippocampus [38,42]. Importantly, the timing, pattern, and extent of Tau phosphorylation and aggregation in *PS19* mice show some sex specificity, suggesting that differences in Tau phosphorylation dynamics between males and females may contribute differently to NFT formation [44].

In this study, we asked whether GDE2 regulates Tau phosphorylation in the *PS19* tauopathy model, which lacks amyloid pathology. We found that loss of GDE2 transiently modulates Tau phosphorylation at select residues in a spatially and sex-specific manner. Moreover, changes in Tau phosphorylation coincided with region- and sex-specific alterations in GSK3α/β and AKT activity, while phosphatase activity was unchanged. These findings put forth GDE2 as a regulator of kinase-mediated Tau phosphorylation independent of amyloid pathology, suggesting its role in tauopathy may be distinct from, and potentially opposed to, its role in amyloid-driven disease contexts. Together, these results further our mechanistic understanding of how upstream signaling influences Tau phosphorylation dynamics in disease and highlight the importance of disease context when evaluating potential regulators of Tau pathology.

## Results

### Tau phosphorylation at select pro-aggregation residues is decreased in the cortex of *PS19*;*Gde2KO* animals

*PS19* animals show increased Tau phosphorylation across different residues over time [38]. To determine whether GDE2 contributes to Tau phosphorylation, we crossed *Gde2* knockout mice with the *PS19* tauopathy model and aged *PS19;Gde2KO* and *PS19* (control) mice to 3, 6, and 9 months. We then directly compared pTau:Tau ratios in cortical extracts from age-matched *PS19* and *PS19;Gde2KO* animals by western blot, focusing on four residues with distinct roles in Tau pathology: S202/T205, a pro-aggregation site that increases sharply at moderate stages of pathology and detected by the AT8 epitope; T212, a pro-aggregation site that increases continuously from early to late stages of disease; S396, a pro-aggregation site considered among the first to become hyperphosphorylated in disease; and S262, a site associated with Tau detachment from microtubules that is linked to early-stage Tau pathology [10,14,24,27–30].

At 3 months, western blot analysis showed no significant differences between genotypes in the pTau:Tau ratios at any of the four residues examined (S202/T205, T212, S396, S262; Fig. S1 A-D) across the total cohort of male and female animals. Because *PS19* mice show sex-specific differences in the onset and extent of Tau phosphorylation, we also analyzed pTau:Tau ratios separately in males and females (Fig. S1 A-D). No significant changes in Tau phosphorylation of the four residues were found; however, we noticed a downward trend in pTau:Tau ratios at S202/T205 (p = 0.058) in male *PS19;Gde2KO* mice compared with *PS19* controls (Fig. S1 A).

At 6 months, pTau(S202/T205):Tau and pTau(T212):Tau ratios were significantly decreased in *PS19;Gde2KO* cortex relative to *PS19* (Fig. 1 A,B) in the total cohort encompassing male and female animals. Interestingly, the reductions in pTau(S202/T205):Tau and pTau (T212) were restricted to males (Fig. 1 A,B). In contrast, pTau:Tau ratios at residues S262 and S396 did not differ significantly between genotypes at 6 months (Fig. S2 A,B). To independently confirm the reduction in the pTau:Tau ratios at residues S202/T205 and T212, we stained cortical sections from 6-month-old *PS19* and *PS19;Gde2KO* animals with antibodies against the neuronal marker NeuN and pTau(S202/T205) or pTau(T212), imaged the entorhinal and piriform cortex, and quantified pTau^+^/NeuN^+^ cells. Consistent with the western blot results, 6-month *PS19;Gde2KO* cortices showed fewer pTau(S202/T205)^+^ neurons compared with controls in the total cohort and in males (Fig. 1 C). While there was a significant reduction in pTau(T212)^+^ neurons in *PS19;Gde2KO* cortices relative to *PS19* within the total cohort (Fig. 1 D), the downward trends observed in male and female animals did not reach significance due to inherent variability in controls. Together, our biochemical and histological data suggest that GDE2 loss selectively decreases Tau phosphorylation at pro-aggregation sites S202/T205 and T212.

**Fig. 1:**
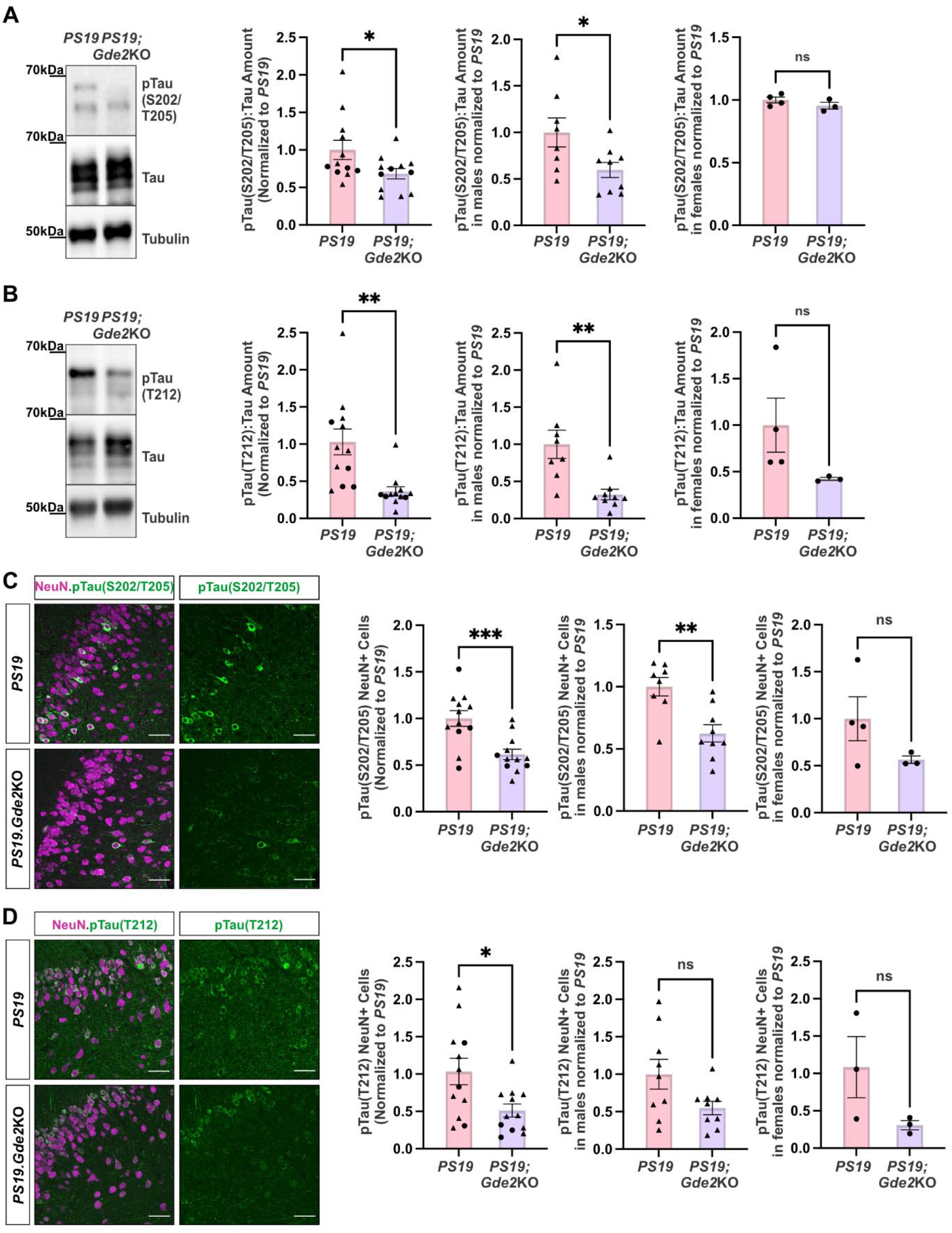
Tau phosphorylation at S202/T205 and T212, is decreased in 6-month *PS19*;*Gde2KO* cortex. **A, B,** Representative western blots of 6-month *PS19* and *PS19;Gde2KO* cortical extracts and graphs with corresponding quantification for the overall cohort (left), males (middle), and females (right). Triangles and dots denote males and females, respectively. **A**, pTau (S202/T205):Tau ratio; left: *p=0.0386, middle: *p=0.0315, right: ns p=0.2559. **B**, pTau(T212):Tau ratio; left: **p=0.0016, middle: **p=0.0032, right: ns p=0.1552. **C**,**D** Immunostaining of pTau(S202/T205) (**C**) and pTau(T212) (**D**) in 6-month *PS19* and *PS19;Gde2KO* entorhinal/piriform cortex with corresponding quantification for the overall cohort (left), males (middle), and females (right). Triangles and dots denote males and females, respectively. **C**, pTau(S202/T205); left: ***p=0.0009, middle: **p=0.0022, right: ns p=0.1784. **D**, pTau(T212); left: *p=0.0150, middle: ns p=0.0557, right: ns p=0.1333. All graphs: *PS19* n=12 (8 males, 4 females), *PS19;Gde2*KO n=12 (9 males, 3 females). Mean ± SEM, two-tailed unpaired Student’s *t*-test. Scale bars: **C**, **D**, 50μm.

To determine whether GDE2 loss continued to affect Tau phosphorylation during late-stage tauopathy, we quantified pTau:Tau ratios at residues S202/T205, T212, S262, and S396 in cortical extracts from 9-month *PS19* and 9-month *PS19;Gde2KO* animals. We observed no significant differences in pTau:Tau ratios at any of these residues between genotypes (Fig. S3 A-D), either in the total cohort of mice or when separated by sex. Taken together, these observations suggest that the effects of GDE2 loss on Tau phosphorylation are transient.

### *Gde2* ablation has no discernible effects on neuronal loss or Tau aggregation

Given the reduced pTau:Tau ratios at S202/T205 and T212 in the 6-month *PS19;Gde2KO* cortex in males, we next asked whether GDE2 ablation affects neuronal loss or Tau aggregation in 6-month *PS19;Gde2KO* males. To measure neuronal loss, we identified neurons by NeuN immunostaining together with the DNA-binding dye Hoechst 33342, and quantified neuronal numbers in the cortex of 6-month- and 9-month-old *PS19* and *PS19;Gde2KO* male mice (see methods for more details). We detected no differences in neuronal numbers between genotypes at either time point (Fig. 2 A,B). To evaluate Tau aggregation, we dissected cortices from 6-month-old *PS19* and *PS19;Gde2KO* male mice and used established sequential extraction methods to enrich for insoluble (RIPA-soluble fraction) and highly insoluble (Formic Acid (FA)-soluble fraction) forms of Tau [38,45,46]. Consistent with our earlier observations, we detected reduced pTau(T212) levels in male *PS19;Gde2KO* cortices compared with *PS19* controls (Fig. 2 C). However, the amounts of insoluble and highly insoluble forms of pTau(T212) were not significantly different between genotypes (Fig. 2 C). Lastly, we evaluated NFT formation via Gallyas silver staining of cortical sections from 9-month-old *PS19* and *PS19;Gde2KO* male mice and found no differences in NFTs between genotypes (Fig. 2 D,E). These observations suggest that the delay in Tau phosphorylation caused by *Gde2* ablation in males does not measurably affect these end-stage pathologies, consistent with a transient rather than sustained effect of GDE2 loss on Tau phosphorylation at S202/T205 and T212.

**Fig. 2:**
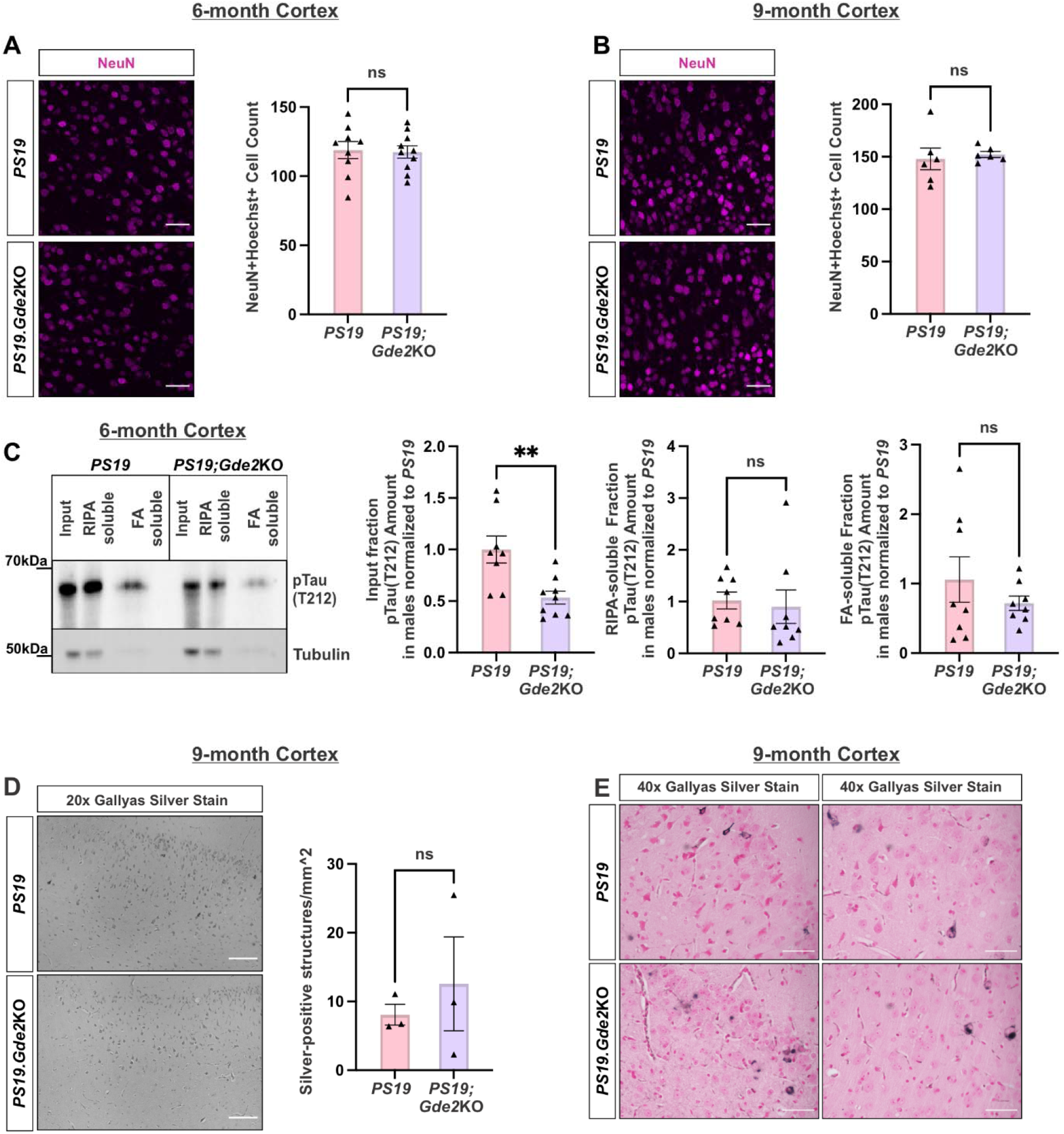
End-stage pathology in males is unchanged between *PS19* and *PS19*;*Gde2KO* cortex. **A**, **B**, Immunostaining of NeuN in male 6-month and 9-month *PS19* and *PS19;Gde2KO* cortex with corresponding quantification. **A**, 6-month, ns p=0.8573, **B**, 9-month, ns p=0.7013. **C**, Representative western blots of pTau(T212) levels in male 6-month *PS19* and *PS19;Gde2KO* following FA fractionation of cortical extracts with corresponding quantification for the input fraction (left: **p=0.0046), RIPA-soluble fraction (middle, ns p= 0.7463), and FA-soluble fraction (right, ns p=0.3389). **D**, Representative 20x images of Gallyas Silver Stain in 9-month *PS19* (top) and *PS19;Gde2KO* (bottom) cortex with corresponding quantification (ns p=0.5565). **E**, Representative 40x images of Gallyas Silver Stain, showing silver-positive structures in 9-month *PS19* (top panels) and *PS19;Gde2KO* (bottom panels) cortex. All graphs: Mean ± SEM, two-tailed unpaired Student’s *t*-test. **A-B**: *PS19*: 6mo n=9, 9mo n=6; *PS19;Gde2KO*: 6mo n=10, 9mo n=6. **D**: *PS19*: n=3; *PS19;Gde2KO* n=3. Scale bars: **A-B, E**, 50μm; **D**, 100μm.

### Phosphorylated Tau at S262 is increased in the hippocampus of *PS19*;*Gde2KO* animals

Tau phosphorylation in disease affects multiple regions in the brain, including the hippocampus [15,39]. To determine whether GDE2 has region-specific effects on Tau phosphorylation, we repeated our analysis, this time focusing on the hippocampus. We quantified pTau:Tau ratios at residues S202/T205, T212, S262, and S396 in hippocampal extracts from *PS19;Gde2KO* and *PS19* control animals at 3, 6, and 9 months by western blot and analyzed pTau:Tau ratios across the entire cohort of animals and separately, in males and females.

At 3 months, none of the four ratios differed significantly between genotypes across the cohort or in males and females (Fig. S4 A-D). However, we noted a downward trend in pTau(T212):Tau (p = 0.053) and pTau(S396):Tau (p = 0.066) ratios across the cohort (Fig. S4 B,D). At 6 months, when pathology is typically moderate, the pTau:Tau ratios at S202/T205, T212, and S396 also did not differ significantly between *PS19* and *PS19;Gde2KO* hippocampus within the cohort or when separated by sex (Fig. S5 A-C). However, the pTau:Tau ratio at S262 was increased in *PS19;Gde2KO* hippocampus relative to *PS19*, and this increase was observed in the total cohort and in both males and females (Fig. 3 A). To confirm the increase in the pTau(S262):Tau ratio observed via western blot, we stained hippocampal sections from 6-month-old *PS19* and *PS19;Gde2KO* animals with pTau(S262) and the neuronal marker NeuN. Males and females were grouped because GDE2 loss showed no sex-specific effects on Tau phosphorylation in hippocampal extracts (Fig. 3 A). Consistent with our western blot data, *PS19;Gde2KO* hippocampus showed an increase in the number of pTau(S262)^+^ neurons relative to *PS19* (Fig. 3 B). Together, these observations suggest that the loss of GDE2 increases Tau phosphorylation specifically at S262 in the hippocampus at 6 months.

**Fig. 3:**
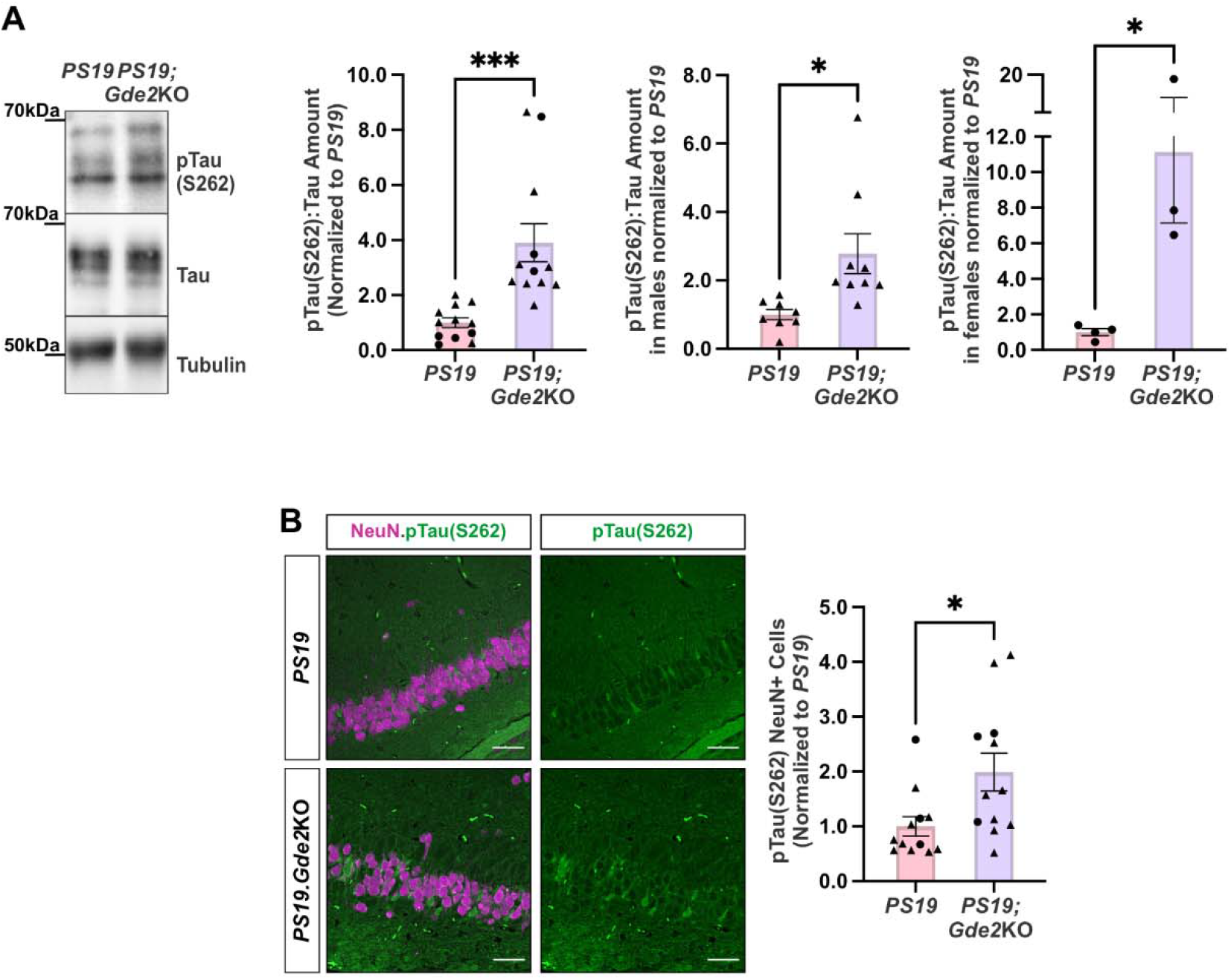
Tau phosphorylation at S262 is increased in 6-month *PS19*;*Gde2KO* hippocampus. **A**, Representative western blot of 6-month *PS19* and *PS19;Gde2KO* hippocampal extracts with corresponding quantification for pTau(S262):Tau ratio for the overall cohort (left, ***p=0.0005), males (middle, *p=0.0136), and females (right, *p=0.0293). Triangles and dots denote males and females, respectively. **B**, Immunostaining of pTau(S262) in 6-month *PS19* and *PS19;Gde2KO* hippocampus with corresponding quantification (*p=0.0181). Triangles and dots denote males and females, respectively. All graphs: *PS19* n=12 (8 males, 4 females), *PS19;Gde2*KO n=12 (9 males, 3 females). Mean ± SEM, two-tailed unpaired Student’s *t*-test.

To determine whether the loss of GDE2 has an impact on Tau phosphorylation during late-stage tauopathy, we compared the pTau:Tau ratios at residues S202/T205, T212, S262, and S396 in hippocampal extracts from 9-month *PS19* and 9-month *PS19;Gde2KO* animals. There were no significant differences in pTau:Tau ratios at all four residues between genotypes in the total cohort or when separated by sex (Fig. S6 A-D), suggesting that the loss of GDE2 has a transient effect on Tau phosphorylation. We then asked whether GDE2 loss affects neuron loss or Tau aggregation in the hippocampus at end-stage disease. Because GDE2 shows no sex-specific effects on Tau phosphorylation in the hippocampus, we grouped both males and females for these analyses. Quantification of NeuN+/Hoechst+ neurons in the hippocampus showed comparable neuron counts between *PS19* and *PS19;Gde2KO* animals at 6 months and 9 months (Fig. 4 A,B), and Gallyas silver staining analyzed across all hippocampal compartments revealed no difference in NFTs between 9-month *PS19* and *PS19;Gde2KO* animals (Fig. 4 C,D). Together, these observations indicate that the transient change in hippocampal Tau phosphorylation caused by *Gde2* ablation does not measurably affect neuronal loss or NFT formation.

**Fig. 4:**
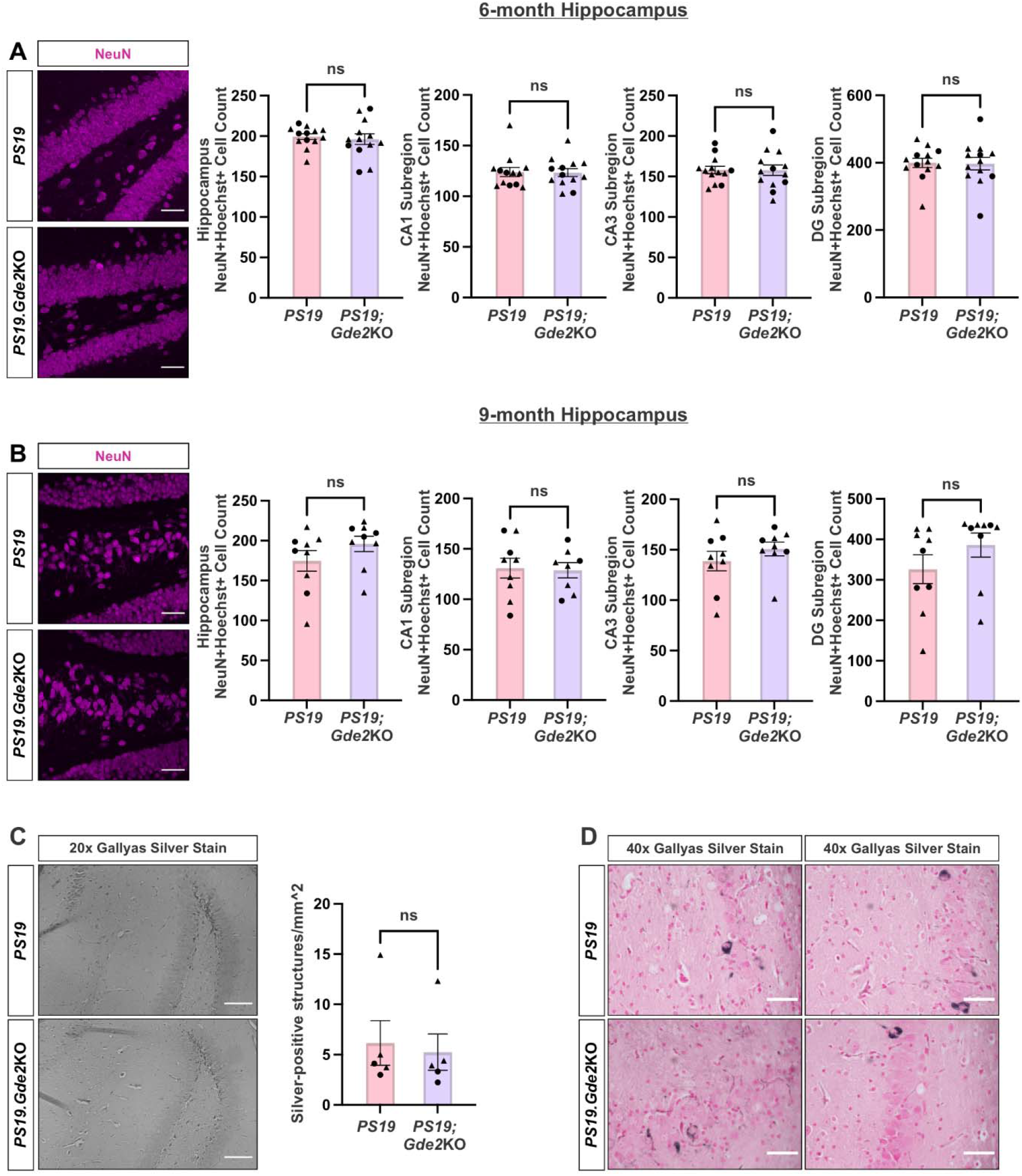
End-stage pathology is unchanged between *PS19* and *PS19*;*Gde2KO* hippocampus. **A-B**, Immunostaining of NeuN in 6-month and 9-month *PS19* and *PS19;Gde2KO* hippocampus with corresponding quantification for the overall hippocampus and the CA1, CA3, and dentate gyrus subregions. Triangles and dots denote males and females, respectively. **A**, 6-month; hippocampus: ns p=0.6816, CA1: ns p=0.9265, CA3: ns p=0.9716, DG ns p=0.9226. **B**, 9-month; hippocampus: ns p=0.2037, CA1: ns p=0.8700, CA3: ns p=0.3241, DG ns p=0.2163. **C**, Representative 20x images of Gallyas Silver Stain in 9-month *PS19* (top) and *PS19;Gde2KO* (bottom) hippocampus with corresponding quantification (ns p=0.7588). Triangles and dots denote males and females, respectively. **D**, Representative 40x images of Gallyas Silver Stain, showing silver-positive structures in 9-month *PS19* (top panels) and *PS19;Gde2KO* (bottom panels) hippocampus. All graphs: Mean ± SEM, two-tailed unpaired Student’s *t*-test. **A-B**: *PS19*: 6mo n=13, 9mo n=9; *PS19;Gde2KO*: 6mo n=13, 9mo n=9. **C**: *PS19* n=5, *PS19;Gde2KO* n=5. Scale bars: **A**,**B**,**D**, 50μm; **C**, 100μm.

### Kinase activity is altered in the cortex and hippocampus of 6-month *PS19*;*Gde2KO* animals

Tau phosphorylation homeostasis reflects a balance between kinase and phosphatase activity, which is disrupted in disease [15,16]. GSK3α/β directly phosphorylates Tau at multiple residues, while AKT modulates Tau phosphorylation both directly and indirectly. The activity of both enzymes is regulated by phosphorylation: GSK3α/β is inhibited by phosphorylation, whereas AKT activity requires phosphorylation [47]. Accordingly, to determine if GDE2 modulates GSK3α/β and AKT kinase activity, we used pGSK3α/β:GSK3α/β and pAKT:AKT ratios as proxies for kinase activity in cortical and hippocampal lysates from 6-month *PS19* and *PS19;Gde2KO* animals. We quantified phosphorylation of GSK3α at S21 and GSK3β at S9 to measure GSK3α/β kinase inhibition, and pAKT at S473 to measure AKT kinase activation. In the cortex, the pGSK3α/β:GSK3α/β ratio was increased in *PS19;Gde2KO* relative to *PS19* across the cohort and in both males and females (Fig. 5 A), consistent with reduced GSK3α/β activity. While pAKT:AKT ratios were reduced in the total cohort and in *PS19;Gde2KO* females, no changes were observed in males implying that AKT kinase activity is unchanged in *PS19;Gde2KO* males (Fig. 5 B). The reduction in cortical GSK3α/β activity aligns with the observed decreases in pTau(S202/T205):Tau and pTau(T212):Tau ratios in the male *PS19;Gde2KO* cortex (Fig. 1 A-D) and accordingly, raises the possibility that GDE2 regulates GSK3α/β activity to modulate Tau phosphorylation at these sites. In the hippocampus, the pGSK3α/β:GSK3α/β ratio did not differ between genotypes across the cohort and in males but was elevated in females in *PS19;Gde2KO* relative to *PS19* indicative of GSK3α/β inhibition (Fig. 5 C). However, the pAKT:AKT ratio was markedly increased in *PS19;Gde2KO* relative to *PS19* in the total cohort and in males and females, consistent with elevated AKT kinase activity in both sexes (Fig. 5 D). The increase in pAKT:AKT in the hippocampus of *PS19;Gde2KO* males and females coincides with the increased S262 phosphorylation we observed in the hippocampus of both sexes (Fig. 3 A,B). These observations suggest that GDE2 may act as an upstream modulator of GSK3α/β and AKT kinase activity, with region-specific effects in the cortex and hippocampus that coincide with Tau phosphorylation at pro-aggregation and dissociation residues, respectively.

**Fig. 5:**
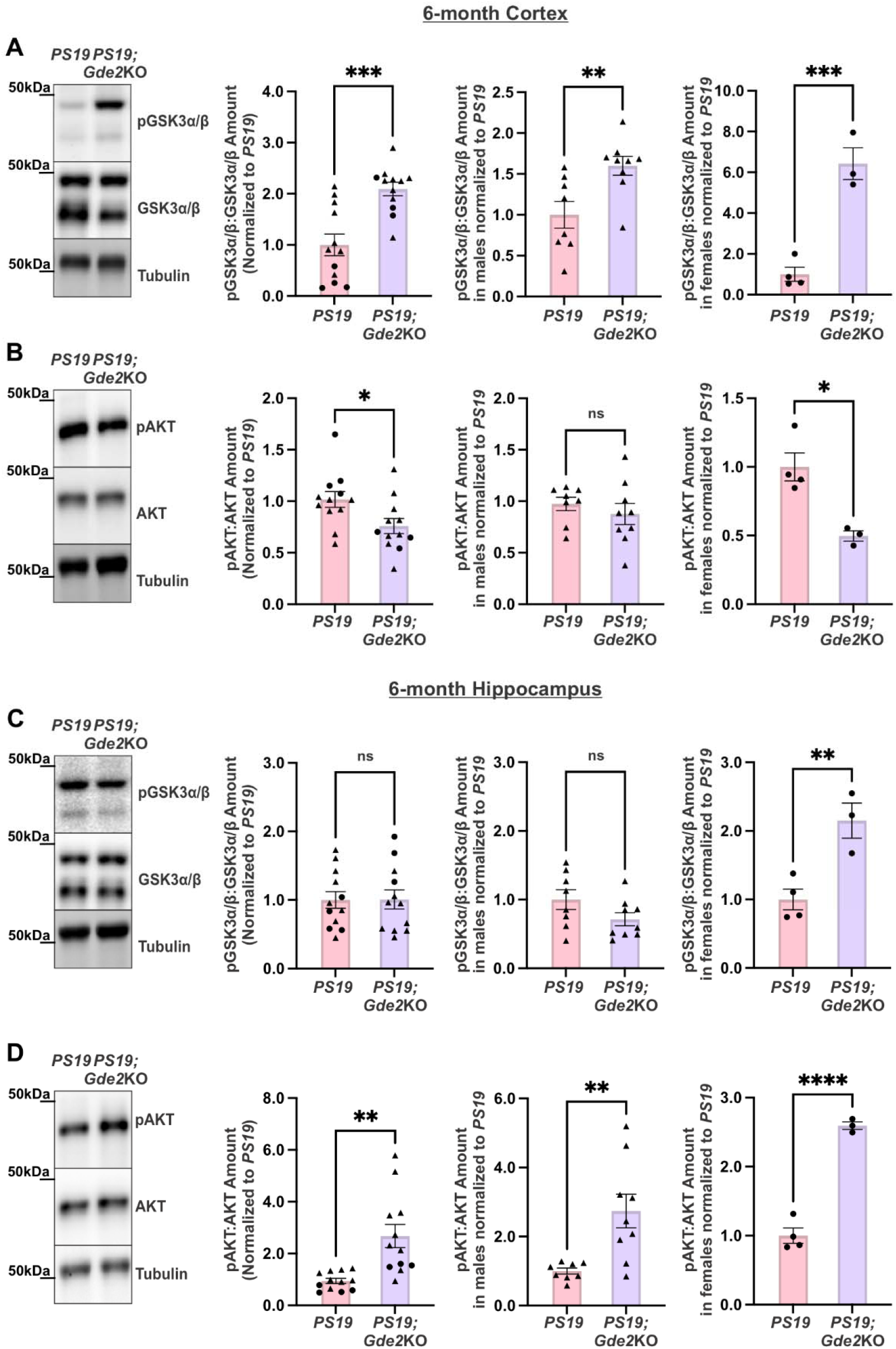
Kinase activity is altered in the cortex and hippocampus of 6-month *PS19*;*Gde2KO* animals. **A-D,** Representative western blots of 6-month *PS19* and *PS19;Gde2KO* cortical (**A**, **B**) and hippocampal (**C**, **D**) extracts with corresponding quantification for the overall cohort (left), males (middle), and females (right). Triangles and dots denote males and females, respectively. **A**, **C**, pGSK3α/β:GSK3α/β ratios in 6-month *PS19* and *PS19;Gde2KO* cortical and hippocampal extracts with corresponding quantification. **A**, cortex; left: ***p=0.0002, middle: **p=0.0080, right: ***p=0.0009. **C**, hippocampus; left: ns p=0.9710, middle: ns p=0.1096, right: **p=0.0091. **B**, **D**, pAKT:AKT ratios in 6-month *PS19* and *PS19;Gde2KO* cortical and hippocampal extracts with corresponding quantification. **B**, cortex; left: *p=0.0243, middle: ns p=0.4498, right: *p=0.0101. **D**, hippocampus; left: **p=0.0010, middle: **p=0.0047, right: ****p<0.0001. All graphs: *PS19* n=12 (8 males, 4 females), *PS19;Gde2*KO n=12 (9 males, 3 females). Mean ± SEM, two-tailed unpaired Student’s *t*-test.

We next asked whether GDE2 also modulates phosphatase activity, focusing on PP1, which acts directly on Tau at specific residues, and PP2A, the main phosphatase acting on Tau [15,21]. As phosphorylation inhibits the activity of both PP1 and PP2A [48,49], we calculated the ratios of phosphorylated to total phosphatase levels as a proxy for phosphatase activity. Western blot analysis of cortical and hippocampal extracts prepared from *PS19* and *PS19;Gde2KO* animals showed no significant changes in pPP1:PP1 and pPP2A:PP2A ratios between genotypes in the total cohort or between sexes (Fig. S7 A-D). Together, these findings suggest that GDE2 does not act through PP1 or PP2A to alter Tau phosphorylation in the context of tauopathy in the cortex or the hippocampus.

### GDE2 expression increases Tau phosphorylation at S202/T205 in vitro

GDE2 is expressed in neurons, subsets of oligodendrocytes, and vascular endothelial cells [31,33]. To determine the contribution of neuronal GDE2 to modulating Tau phosphorylation, we cultured primary cortical neurons from postnatal day (P)0 *PS19* and *PS19;Gde2KO* pups and harvested them after 21 days in vitro (DIV). We focused on Tau phosphorylation at S202/T205 because it is among the most prominently phosphorylated residues in disease and showed reduced phosphorylation in *PS19;Gde2KO* cortex in vivo [11–13]. Western blot analysis showed a decrease in the pTau(S202/T205):Tau ratio in *PS19;Gde2KO* DIV21 primary cortical neurons compared with *PS19* controls (Fig. 6 A), suggesting that neuronal GDE2 modulates Tau phosphorylation in neurons in a cell-autonomous manner.

**Fig. 6:**
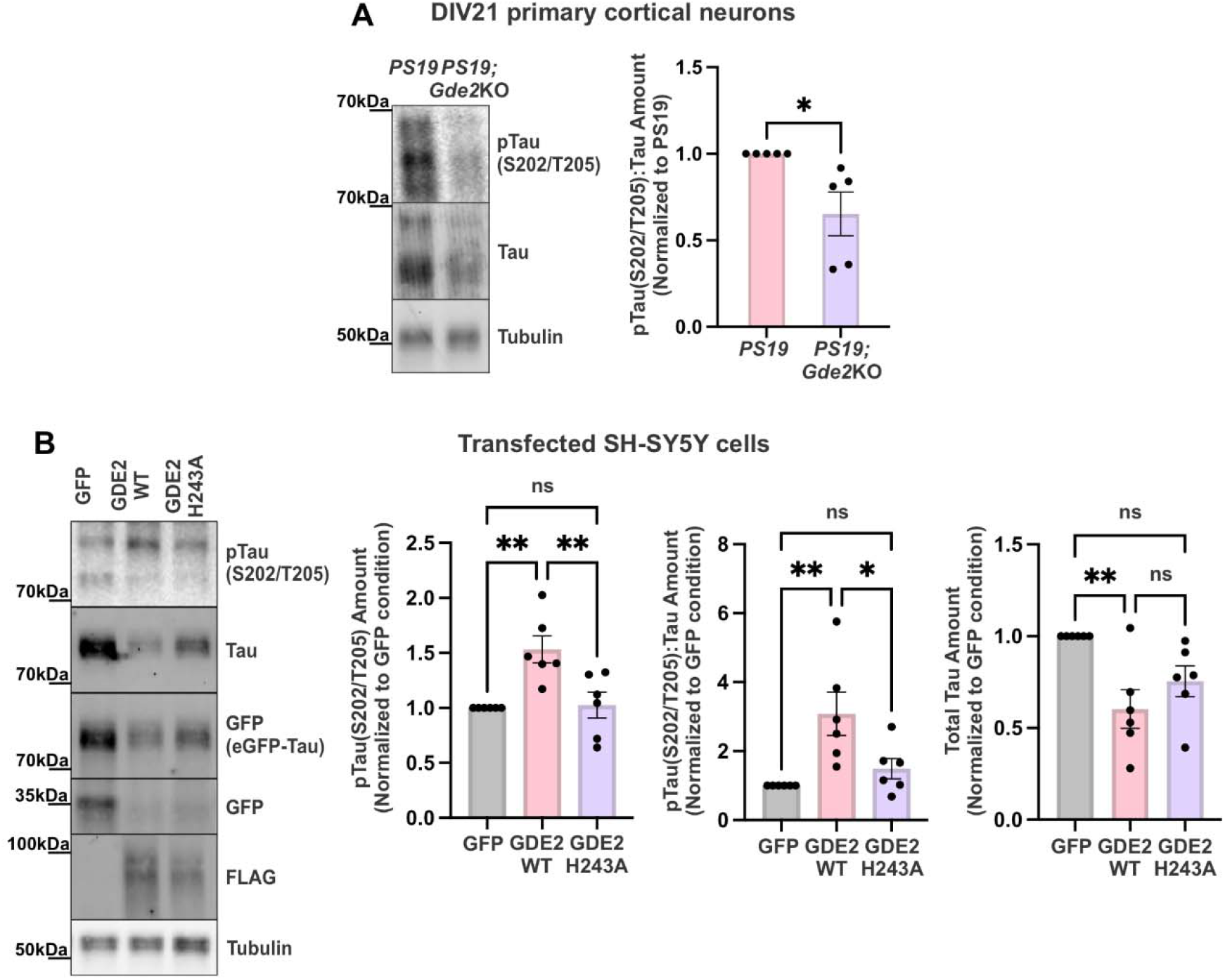
GDE2 expression results in increased tau phosphorylation at S202/T205 *in vitro*. **A**, Representative western blot and quantification for pTau(S202/T205):Tau ratio in DIV 21 primary cortical neurons (*p=0.0250). **B**, Representative western blot and quantification for pTau(S202/T205), pTau(S202/T205):Tau ratio, and Total Tau in transfected SH-SY5Y cells (left: **p=0.0035 (GFP vs GDE2 WT), **p=0.0050 (GDE2 WT vs GDE2H243A), ns p=0.9811 (GFP vs GDE2H243A); middle: **p=0.0057 (GFP vs GDE2 WT), *p=0.0319 (GDE2 WT vs GDE2H243A), ns p=0.6665 (GFP vs GDE2H243A); right: **p=0.0066 (GFP vs GDE2 WT), ns=0.3735 (GDE2 WT vs GDE2H243A), ns p=0.0967 (GFP vs GDE2H243A)). **A**: *PS19* n=5, *PS19;Gde2KO* n=5; **B**: GFP n=6, GDE2 WT n=6, GDE2H243A n=6. All graphs: Mean ± SEM. Each symbol represents one biological replicate (see methods for normalization information). **A**: two-tailed unpaired Student’s *t*-test, **B**: one-way ANOVA with Tukey’s multiple comparisons test.

GDE2 is a six-transmembrane protein with an extracellular glycerophosphodiester phosphodiesterase (GDPD) catalytic domain that cleaves the GPI-anchor that tethers select proteins to the membrane [31,32]. Previous studies have shown that a single-point mutation in the GDPD domain (GDE2-H243A) abolishes GDE2 GPI-anchor-cleaving activity without compromising its surface expression [50]. To test whether GDE2 GPI-anchor cleaving activity is required to modulate Tau phosphorylation, we co-transfected plasmids expressing eGFP-Tau with either plasmids expressing GFP, FLAG-tagged GDE2-WT, or FLAG-tagged GDE2-H243A into undifferentiated SH-SY5Y cells. We then harvested cells 3 days post-transfection, and subjected cell lysates to western blotting. GDE2-WT expression led to an increase in pTau(S202/T205) levels and pTau(S202/T205):Tau ratio, whereas catalytically dead GDE2-H243A had no significant effect on either measure (Fig. 6 B). Interestingly, GDE2-WT, but not GDE2-H243A, expression led to a decrease in total Tau levels relative to the GFP control condition (Fig. 6 B). Together, these observations suggest that GDE2 modulation of Tau phosphorylation and total Tau levels is dependent on its GPI-anchor-cleaving function.

## Discussion

Here, we find that the six-transmembrane GPI-anchor-cleaving enzyme, GDE2, modulates Tau phosphorylation in a region- and sex-specific manner in the context of the *PS19* mouse tauopathy model. Loss of GDE2 in *PS19* mice at 6 months of age changes Tau phosphorylation at select residues, with reduced pTau:Tau ratios at S202/T205 and T212 in the cortex of male mice, and increased pTau:Tau ratio at S262 in the hippocampus in male and female animals. These phosphorylation changes coincide with changes in kinase, but not phosphatase, activity: GSK3α/β kinase activity was decreased at 6 months in male *PS19;Gde2KO* cortex, while AKT kinase activity was increased in the 6-month hippocampus of male and female *PS19;Gde2KO* animals. Analysis of *PS19;Gde2KO* cultured primary cortical neurons and mechanistic studies in SH-SY5Y cells indicate, respectively, that GDE2 regulation of Tau phosphorylation at S202/T205 is cell-autonomous and requires its GPI-anchor-cleaving function. These observations suggest possible roles for GDE2 in the complex regulation of kinase-dependent Tau phosphorylation in the context of tauopathy, with region-specific modulation of Tau phosphorylation at select residues associated with pro- and anti-aggregation properties.

Previous studies have shown that GDE2 cleaves and inactivates the GPI-anchored metalloprotease inhibitor REversion-inducing Cysteine-rich protein with Kazal motifs (RECK), thereby promoting the neuroprotective, non-amyloidogenic cleavage of Amyloid Precursor Protein (APP) by the α-secretase A Disintegrin And Metalloproteinase Domain 10 (ADAM10) [34]. Consistent with this function, loss of GDE2 in the *APP/PS1* mouse model of amyloidosis accelerates plaque deposition [34]. Notably, loss of GDE2 in *APP/PS1* mice also increased Tau phosphorylation at the pro-aggregation site S202/T205 [34]. These observations suggest that loss of GDE2 in this amyloidosis model contributes to amyloid pathology and promotes aggregation-prone Tau phosphorylation. In contrast, our current study indicates that, in the context of tauopathy alone, loss of GDE2 appears to be neuroprotective, as it reduces Tau phosphorylation at pro-aggregation sites and increases phosphorylation at S262, which promotes Tau dissociation from microtubules. Thus, depending on the context, loss of GDE2 can modulate Tau phosphorylation towards states that favor or mitigate Tau aggregation. Although the mechanisms underlying these contrasting effects require further study, our findings highlight the complexity of disease-relevant pathways and underscore the need to study them in more disease-relevant contexts.

We find that GDE2 ablation in *PS19* animals does not uniformly alter Tau phosphorylation across the cortex and hippocampus: Tau phosphorylation at S202/T205 and T212 decreases in the cortex, while Tau phosphorylation at S262 increases in the hippocampus. Interestingly, these changes are sex-specific: males show cortical reductions in pTau at S202/T205 and T212, while males and females show hippocampal increases in pTau at S262. The mechanisms underlying these sex-specific differences remain unclear. Nevertheless, the region-specific effects of GDE2 loss in the *PS19* model align with mitigating Tau aggregation rather than with blanket effects that promote or reduce Tau phosphorylation. The different cortical and hippocampal changes in Tau phosphorylation observed in response to GDE2 loss coincide with sex- and region-specific changes in kinase but not phosphatase activity, suggesting that GDE2 regulates the local kinase environment to modulate Tau phosphorylation. In the cortex, GSK3α/β activity was decreased in *PS19;Gde2KO* animals, while AKT activity was increased in the hippocampus, suggesting that these perturbations might underlie the changes in Tau phosphorylation observed in *PS19;Gde2KO* animals. Interestingly, female *PS19;Gde2KO* mice showed increased phosphorylation of GSK3α/β and AKT in the hippocampus, consistent with activation of the canonical AKT-GSK3α/β pathway, in which AKT directly phosphorylates GSK3α/β to inhibit its activity [51]. This is consistent with prior observations in *Gde2KO* mice that suggest physiological roles for GDE2 in regulating the canonical AKT-GSK3α/β pathway in the hippocampus [51]. In contrast, the relationship between GSK3α/β and AKT phosphorylation indicative of the canonical pathway is not observed in male or female *PS19;Gde2KO* cortex or in male *PS19;Gde2KO* hippocampus, suggesting that GDE2 acts on AKT and GSK3α/β independently of the canonical AKT-GSK3α/β pathway in these structures. Together, these observations further support that GDE2 functions in a context-dependent and region-specific manner in the brain.

Our analysis of GDE2-mediated modulation of Tau phosphorylation suggests a transient role for GDE2 in regulating Tau phosphorylation rather than the continual maintenance of Tau phosphorylation homeostasis in *PS19* animals. Indeed, GDE2 ablation does not alleviate Tau aggregation, NFT formation, or neuronal loss in *PS19* animals at 6 and 9 months, suggesting that the delay and transient reductions in Tau phosphorylation resulting from GDE2 ablation are not sufficient to alter end-stage pathology. One possible reason is that GDE2 loss affects only a subset of residues, whose effects may be obscured in the *PS19* model, which is an overexpression model prone to robust hyperphosphorylation [38]. Alternatively, the impact of GDE2 loss on S202/T205 and T212 phosphorylation in the cortex and S262 phosphorylation in the hippocampus in this overexpression model may not substantially affect Tau’s propensity to aggregate in these brain areas. Further, it is possible that pre-fibrillar Tau resulting from GDE2-dependent effects on Tau phosphorylation may be rapidly cleared before any changes to Tau aggregation are evident.

GDE2 is expressed in neurons, vascular endothelial cells, and a subset of terminally differentiated oligodendrocytes [33]. The observed decrease in the pTau(S202/T205):Tau ratio in *PS19;Gde2KO* primary cortical neurons suggests roles for neuronally expressed GDE2 in modulating Tau phosphorylation at this residue in a cell-autonomous manner. Whether neuronal GDE2 also regulates Tau phosphorylation at T212 and S262 remains an open question because of limitations in growing primary cortical and hippocampal neurons in vitro for extended periods. Future studies using cell-type-specific genetic ablation of GDE2 in vivo would be a feasible approach to define the cell-type-specific requirements for GDE2 in modulating Tau phosphorylation at different residues over time. Our studies in SH-SY5Y cells provide strong evidence that GDE2 acts via its GPI-anchor-cleaving enzymatic function and indicate that GDE2-dependent regulation of a target GPI-anchored protein(s) is required for its effects on Tau phosphorylation. The identity of the GPI-anchored target remains to be elucidated. However, potential candidates include the metalloprotease inhibitor RECK and the heparan sulfate proteoglycan GPC6, which drive amyloid and TDP-43 pathology [34,52,53], respectively, when GDE2 function is disrupted, and have been associated with AD and ALS in genome-wide association studies (GWAS) [54–56].

Our study has focused primarily on evaluating GDE2-dependent effects on Tau phosphorylation in *PS19* animals. While a useful model of tauopathy, these animals have several limitations, including overexpression of human Tau (P301S) at levels well above endogenous Tau and a failure to capture the full complexity of human disease [38, 57]. GDE2 is aberrantly localized in intracellular compartments in the postmortem brain of patients with ALS and AD, and biochemical studies support its dysfunction in disease [33,34]. Future studies examining whether GDE2 mislocalization correlates with Tau phosphorylation in postmortem human tissue will be essential to deepen understanding of GDE2’s contributions to Tau pathologies in disease.

## Methods and Materials

### Animal Husbandry

Animals were bred and maintained in accordance with approved Johns Hopkins University Institutional Animal Care and Use Committee protocols. B6;C3-Tg(Prnp-MAPT*P301S)PS19Vle/J (*PS19*) mice were obtained from Jackson Laboratory (Strain#: 008169) and maintained as a hemizygous line. *PS19* animals were crossed with *Gde2KO* animals to generate *PS19;Gde2Het* animals, which were viable and fertile. These were then bred to produce the experimental *PS19* and *PS19;Gde2KO* animals. Animals were genotyped as specified by Jackson Laboratory and as described in *Sabharwal et al., 2011* [58]. Animals were sacrificed within 1 week of the specified 3-, 6-, and 9-month timepoints.

### Paraffin tissue preparation

Mice were anesthetized with 0.02□ml/g Avertin solution (1.3% 2,2,2-Tribromoethanol and 0.7% 2-methyl-2-butanol) in phosphate buffered saline (PBS). Mice were sacrificed via decapitation and brains were dissected. One hemisphere was post-fixed in 4% PFA for 20-24□h, washed with PBS, and prepared for embedding in paraffin blocks. The cortex and hippocampus were isolated from the remaining hemisphere and flash-frozen in an ethanol bath and subsequently processed for biochemical studies.

### Tissue Immunofluorescence

Tissue immunofluorescence was generally performed as described in *Nakamura et al., 2021* [34], with minor modifications. Briefly, tissue was embedded in paraffin, sliced at 4μm, and three to four slices of tissue were mounted per slide. Once dry, slides were placed on a slide warmer for 20 minutes to melt the paraffin wax. Sections were then deparaffinized in Xylenes and rehydrated via serial washes in 100%, 95%, and 70% ethanol with one rinse in distilled water. Tissue was permeabilized with PBS-T (PBS containing 0.3% Triton-X-100) followed by antigen retrieval using sodium citrate buffer (10mM sodium citrate containing 0.5% Tween-20, pH 6.0) for 20 minutes in a 95°C water bath. Sections were then washed in PBS-T and incubated in blocking buffer (5% Bovine Serum Albumin (BSA) in PBS) for 1 hour at room temperature. Sections were then incubated with primary antibodies (diluted in 1% BSA in PBS-T) overnight at 4°C. Sections were washed in PBS before incubating with secondary antibodies (diluted in 1% BSA in PBS) at room temperature for 1 hour. Finally, sections were washed with PBS and then coverslipped using ProLong Diamond Antifade Mountant. Z-stack images were acquired with a Zeiss LSM 700 microscope. At least three images were acquired per tissue slice on each slide for each of the regions analyzed. Specifically, for cortical neuron counts, layers II/III of the entorhinal cortex and layers II/III of the piriform cortex were imaged as these regions of the cortex are known to be among the earliest sites of tau phosphorylation accumulation and also exhibit particular vulnerability to tau pathology [38,59,60]. Neuronal counts were obtained for the CA1, CA3, and dentate gyrus subregions of the hippocampus, and counts across all subregions were averaged to obtain an overall hippocampal neuron count. The same imaging settings were used for all images taken within the same experiment.

### Gallyas Silver Stain

The Gallyas silver stain was performed as described previously in *Li et al., 2016* [61]. Briefly, tissue was embedded in paraffin, sliced at 10μm, and three slices of tissue were mounted per slide. Once dry, slides were placed on a slide warmer for 20 minutes to melt the paraffin wax. Sections were then deparaffinized in Xylenes and rehydrated via serial washes in 100%, 95%, and 70% ethanol with one rinse in distilled water. Sections were incubated in 5% periodic acid for 5 minutes, washed in water, followed by a 1-minute incubation in silver iodide solution (4% sodium hydroxide, 10% potassium iodide, 3.5% of 1% silver nitrate). Sections were then washed with 0.5% acetic acid. Sections were then incubated in developer until the tissue turned light brown. Developer solution is a mix of Solution A (5% anhydrous sodium carbonate in water), Solution B (0.2% ammonium nitrate, 0.2% silver nitrate, and 1% tungstosilic acid in water), and Solution C (0.2% ammonium nitrate, 0.2% silver nitrate, 1% tungstosilic acid, and 0.28% formaldehyde in water), which were mixed at the following ratio: 10:3:7, respectively. Quenching was done with 0.5% acetic acid for 3 minutes, followed by washing with water for 10 minutes. Sections were then incubated in 0.1% gold chloride for 5 minutes to stabilize and better visualize the silver stain. Sections were then washed in water and incubated in 1% sodium thiosulfate for 5 minutes to remove unreacted, residual silver ions. Sections were then washed with tap water and counterstained with 0.1% nuclear fast red for 2 minutes. Finally, sections were washed with tap water, dehydrated via serial washes in 70%, 95%, and 100% ethanol followed by Xylenes, and coverslipped using Sub-X Mounting Media. Images were acquired with a Keyence BZ-X710 epifluorescence microscope. Whole sections were imaged and then stitched together using BZ-X Analyzer software. Silver-positive structures were manually counted and then normalized to area (cells/mm^2^) across the entorhinal and piriform cortices, since these regions of the cortex are known to be among the earliest sites of tau phosphorylation accumulation and exhibit particular vulnerability to tau pathology [38,59,60]. Silver-positive structures were also manually counted across the whole hippocampus and then normalized to area (cells/mm^2^). 40x representative images highlighting the presence of silver-positive structures found in 9-month animals were acquired with a Nikon Eclipse E800 epifluorescence microscope. The same imaging settings were used for all images taken within the same experiment.

### Cell culture

#### Primary cortical neurons

Mouse primary cortical culture was performed as described previously in *Nakamura et al., 2021* [34], with minor modifications. Briefly, cortices were dissected from P0 pups, and cortical tissue was dissociated using 2.5% trypsin for 20 minutes at 37°C. Tissue was washed in plating media (Neurobasal medium supplemented with 10% fetal bovine serum (FBS), 0.5% glucose, 1% sodium pyruvate, and 1% Pen/Strep) and then triturated using a P1000 pipette. Single-cell suspension was achieved by passing the triturated tissue through a 70μm cell strainer. Cells were counted on a hemocytometer and seeded at 125,000 cells/cm^2^ on poly-L-lysine-coated plates. Cells were maintained in maintenance media (Neurobasal medium supplemented with 2% B27 plus, 1% L-glutamine, and 1% Pen/Strep). 5μM cytosine arabinoside was added for 24 hours on DIV3 to inhibit glial growth. From DIV4 onwards, cultures were fed every 3 days and maintained at 37°C with 5% CO_2_ until harvested on DIV21. Each experiment was conducted on the same multi-well plate, with each plate considered a biological replicate. Each biological replicate consists of at least three technical replicates for each group. All replicates from the same experiment were run on the same gel for western blot analysis, and values for the technical replicates of each group were averaged and then normalized to the average of the control group (*PS19*, [62,63]).

#### SH-SY5Y cells

SH-SY5Y cells were purchased from the American Type Culture Collection (ATCC) under the catalog number CRL-2266 (RRID: CVCL_0019). Cells were split at 75% confluency and plated at a 1:50 ratio onto 12-well plates. Cells were maintained in DMEM/F-12 (1:1) medium supplemented with 10% FBS and 1% Pen/Strep. Cultures were fed every 3 days and maintained at 37°C with 5% CO_2_ until harvested. Once cells reached 60% confluency, maintenance media was replaced with antibiotic-free maintenance media overnight before being transfected with eGFP-Tau (pRK5-EGFP-Tau was a gift from Karen Ashe and was acquired from Addgene under plasmid number 4690*4; Hoover et al., 2010,* [64]) and GFP, GDE2, or GDE2-H243A (described previously in *Park et al., 2013* and *Shuler et al., 2024,* [32, 50]) using the Lipofectamine 3000 kit following the manufacturer’s protocol (*ThermoFisher Scientific*). Briefly, 250ng of each plasmid was combined with P3000 Reagent and Opti-MEM. In a separate tube, Lipofectamine and Opti-MEM were combined. These two mixes were incubated at 1:1 for 15 minutes before being applied directly to cells. One hour following transfection, DMEM/F-12 (1:1) medium supplemented with 10% FBS and 1% Pen/Strep was added to transfected wells such that the final concentration of Pen/Strep was 0.5%. Cells were harvested 3 days following transfection. Each experiment was conducted on the same multi-well plate, with each plate considered a biological replicate. Each biological replicate consists of at least three technical replicates for each group. All replicates from the same experiment were run on the same gel for western blot analysis, and values for the technical replicates of each group were averaged and then normalized to the average of the control group (GFP, [62,63]).

### Lysate Preparation

Frozen tissue samples were sonicated in 1mL (cortex) or 500μL (hippocampus) of reassembly buffer containing protease and phosphatase cocktail inhibitor, followed by centrifugation at 50,000xg for 40 minutes at 4°C. 4x Laemmli buffer was added to the supernatant to create a final 1x concentration.

To prepare cell lysates, cells were washed in PBS, lysed directly in 1x Laemmli buffer, sonicated, and centrifuged at 21,000xg for 20 minutes at room temperature.

### Formic Acid Fractionation

Insoluble proteins were obtained through sequential extraction as described in *Ishihara et al., 1999,* [45] and *Julien et al., 2012* [46]. Briefly, frozen cortical and hippocampal samples were sonicated in reassembly buffer containing protease and phosphatase cocktail inhibitor as described above, followed by centrifugation at 50,000xg for 40 minutes at 4°C. The resulting supernatant was diluted with 4x Laemmli buffer to create a 1x concentration before being further diluted 1:18 with 1x Laemmli buffer to create the input fraction. To remove myelin and associated lipids from the samples, the resulting pellet was then homogenized in 1mL (cortex) or 500μL (hippocampus) of 1M sucrose solution in reassembly buffer containing protease and phosphatase cocktail inhibitor using a 22G 1 ½ needle and centrifuged at 50,000xg for 20 minutes at 4°C. The supernatant was discarded and the resulting pellet was dissolved in 600μL (cortex) or 300μL (hippocampus) of RIPA buffer (50mM Tris [pH 8.0], 150 mM NaCl, 0.1% SDS, 0.5% sodium deoxy-cholate, 1% NP40, 5mM EDTA [pH 8.0]) containing protease and phosphatase cocktail inhibitor. Following centrifugation at 50,000xg for 20 minutes at 4°C, the resulting supernatant was diluted to a final 1x concentration using 4x Laemmli buffer and used as the RIPA-soluble fraction. To obtain the insoluble, formic acid (FA) fraction, the resulting pellet was then dissolved and homogenized in 100μL (cortex) or 50μL (hippocampus) of 70% FA. Once homogenized, the FA fraction was neutralized with neutralization buffer (3M Tris, 0.5M NaH_2_PO_4_) at 1:7. 4x Laemmli buffer was then added to create a final 1x concentration.

### Western Blot

Samples were boiled and run on lab-made 10% tris/glycine polyacrylamide gels in buffer before being transferred onto low-fluorescence polyvinylidene difluoride membranes at 100V for 70min at 4°C. Membranes were then blocked with EveryBlot Blocking Buffer for 1 hour at room temperature before applying primary antibodies (diluted in either EveryBlot Blocking Buffer or 2% BSA in TBS-T (tris buffered saline (TBS) containing 0.3% Tween-20)) overnight at 4°C. Membranes were then washed with TBS-T, and either fluorescent protein-conjugated or horseradish peroxidase-conjugated secondary antibodies (diluted in EveryBlot Blocking Buffer) were applied for 1 hour at room temperature. After washing with TBS-T, if needed, membranes were developed with enhanced chemiluminescence substrate. Images were acquired using a ChemiDoc MP imager. Blots were analyzed using ImageJ.

### Statistical analysis

GraphPad Prism (version 11) was used to analyze and plot all data. Pairwise comparisons were calculated using a two-tailed Student’s *t*-test, and multiple comparisons were calculated using a one-way ANOVA with Tukey’s multiple comparisons test. In figures, all values are reported as mean ± SEM (standard error of the mean), all data points represent biological replicates, and asterisks were used to indicate statistical significance as follows: *p<0.05, **p<0.01, ***p<0.001, ****p<0.0001. Specifics for each experiment are included in figure legends.

### Antibodies

Abbreviations used

(MC): monoclonal
(PC): polyclonal
(ms): mouse
(rb): rabbit
(gt): goat
(gp): guinea pig
(dk): donkey
(WB): western blot
(IHC): immunohistochemistry

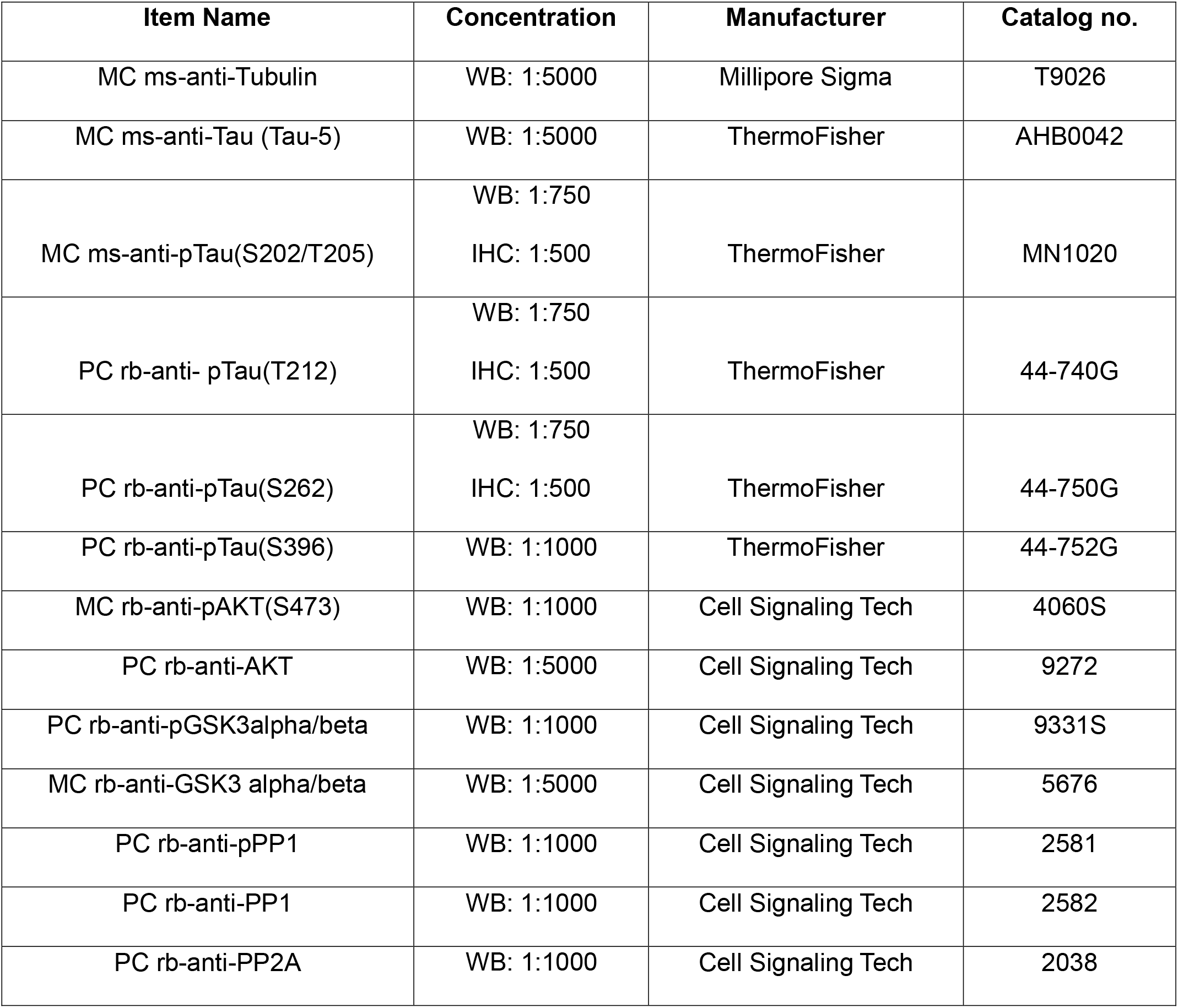
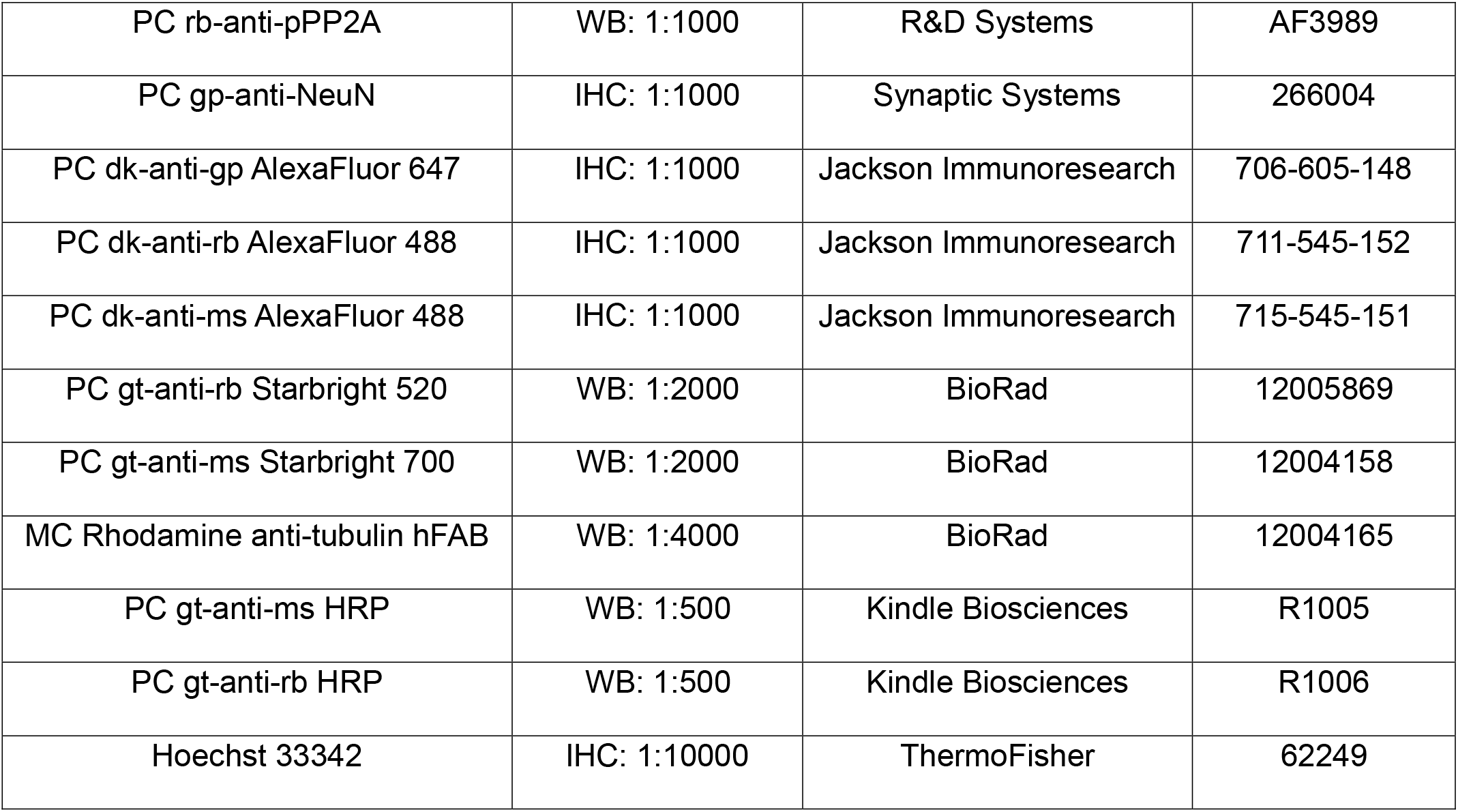

| Item Name | Concentration | Manufacturer | Catalog no. |
|---|---|---|---|
| MC ms-anti-Tubulin | WB: 1:5000 | Millipore Sigma | T9026 |
| MC ms-anti-Tau (Tau-5) | WB: 1:5000 | ThermoFisher | AHB0042 |
| MC ms-anti-pTau(S202/T205) | WB: 1:750<br>IHC: 1:500 | ThermoFisher | MN1020 |
| PC rb-anti- pTau(T212) | WB: 1:750<br>IHC: 1:500 | ThermoFisher | 44-740G |
| PC rb-anti-pTau(S262) | WB: 1:750<br>IHC: 1:500 | ThermoFisher | 44-750G |
| PC rb-anti-pTau(S396) | WB: 1:1000 | ThermoFisher | 44-752G |
| MC rb-anti-pAKT(S473) | WB: 1:1000 | Cell Signaling Tech | 4060S |
| PC rb-anti-AKT | WB: 1:5000 | Cell Signaling Tech | 9272 |
| PC rb-anti-pGSK3alpha/beta | WB: 1:1000 | Cell Signaling Tech | 9331S |
| MC rb-anti-GSK3 alpha/beta | WB: 1:5000 | Cell Signaling Tech | 5676 |
| PC rb-anti-pPP1 | WB: 1:1000 | Cell Signaling Tech | 2581 |
| PC rb-anti-PP1 | WB: 1:1000 | Cell Signaling Tech | 2582 |
| PC rb-anti-PP2A | WB: 1:1000 | Cell Signaling Tech | 2038 |
| PC rb-anti-pPP2A | WB: 1:1000 | R&D Systems | AF3989 |
| PC gp-anti-NeuN | IHC: 1:1000 | Synaptic Systems | 266004 |
| PC dk-anti-gp AlexaFluor 647 | IHC: 1:1000 | Jackson Immunoresearch | 706-605-148 |
| PC dk-anti-rb AlexaFluor 488 | IHC: 1:1000 | Jackson Immunoresearch | 711-545-152 |
| PC dk-anti-ms AlexaFluor 488 | IHC: 1:1000 | Jackson Immunoresearch | 715-545-151 |
| PC gt-anti-rb Starbright 520 | WB: 1:2000 | BioRad | 12005869 |
| PC gt-anti-ms Starbright 700 | WB: 1:2000 | BioRad | 12004158 |
| MC Rhodamine anti-tubulin hFAB | WB: 1:4000 | BioRad | 12004165 |
| PC gt-anti-ms HRP | WB: 1:500 | Kindle Biosciences | R1005 |
| PC gt-anti-rb HRP | WB: 1:500 | Kindle Biosciences | R1006 |
| Hoechst 33342 | IHC: 1:10000 | ThermoFisher | 62249 |

### Other Materials

| Item Name | Manufacturer | Catalog no. |
|---|---|---|
| ProLong Diamond Antifade Mountant | ThermoFisher | P36970 |
| Sub-X Mounting Media | Leica | 3801740 |
| Kwik Quant Kit | Kindle Biosciences | R1004 |
| EveryBlot Blocking Buffer | BioRad | 12010020 |
| Low Fluorescence PVDF Membrane | BioRad | 1620264 |
| Reassembly Buffer | Cepham Life Sciences | 10402-1 |
| Protease+phosphatase inhibitor cocktail | Sigma-Aldrich | PPC1010 |
| Gold chloride solution (200mg/dL) | Sigma-Aldrich | HT1004 |
| Neurobasal medium | ThermoFisher | 21103049 |
| DMEM/F12 (1:1) medium | ThermoFisher | 11320082 |
| Opti-MEM Reduced Serum Medium | ThermoFisher | 31985062 |
| Pen/Strep | ThermoFisher | 15140122 |
| Heat-inactivated FBS | Sigma-Aldrich | F4135-500 |
| B-27 Plus | ThermoFisher | A3582801 |
| L-glutamine (200mM) | ThermoFisher | 25030081 |
| Lipofectamine 3000 Transfection Kit | ThermoFisher | L3000015 |

## Acknowledgements

The authors thank Y. Li for technical assistance; Dr. P. Wong, G. Burns, and M. S. Baghel for help with the Gallyas silver stain protocol; P. Banarjee, J. Llamas Rodriguez, and D. Sama-Borbon for comments on the manuscript.

## Authors and Affiliations

The Solomon Snyder Department of Neuroscience, The Johns Hopkins School of Medicine, 600 N Wolfe St, 1729 Bldg., North, Level 7, Ste. 7305, Baltimore, MD, 21287, USA

Consuelo Jimenez-Ornelas & Shanthini Sockanathan

## Contributions

C.J.O and S.S. conceived the study, designed experiments, interpreted data, and wrote the manuscript. C.J.O. performed the experiments and formal analysis.

## Corresponding Author

Correspondence to Shanthini Sockanathan.

