## Supplemental Figures for "Region-dependent regulation of Tau phosphorylation by Glycerophosphodiester Phosphodiesterase 2 in a mouse model of tauopathy"

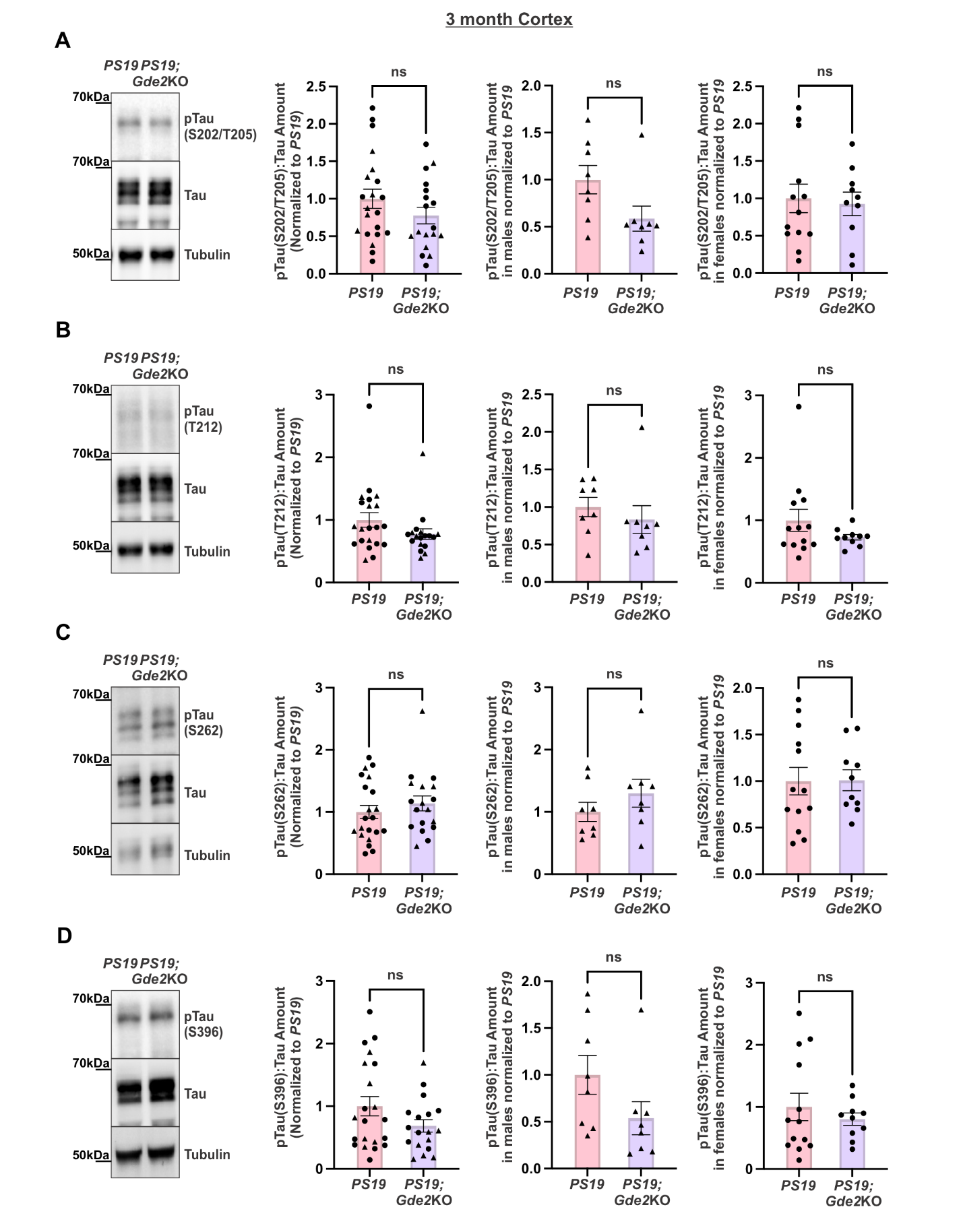
**Fig. S1: Tau phosphorylation is unchanged in the cortex of 3-month *PS19;Gde2*KO animals.** **A-D:** Representative western blots of 3-month *PS19* and *PS19;Gde2KO* cortical extracts and graphs with corresponding quantification for the overall cohort (left), males (middle), and females (right). Triangles and dots denote males and females, respectively. **A**, pTau(S202/T205):Tau ratio; left: ns p=0.1999, middle: ns p=0.0584, right: ns p=0.7774. **B**, pTau(T212):Tau ratio; left: ns p=0.1395, middle: ns p=0.4689, right: ns p=0.2042. **C**, pTau(S262):Tau ratio; left: ns p=0.3888, middle: ns p=0.2889, right: ns p=0.9607. **D**, pTau(S396):Tau ratio; left: ns p=0.1063, middle: ns p=0.1119, right: ns p=0.4705. All graphs: *PS19:* 3mo n=21 (8 males, 13 females); *PS19;Gde2KO*: 3mo n=18 (8 males, 10 females). Mean ± SEM, two-tailed unpaired Student’s *t*-test.

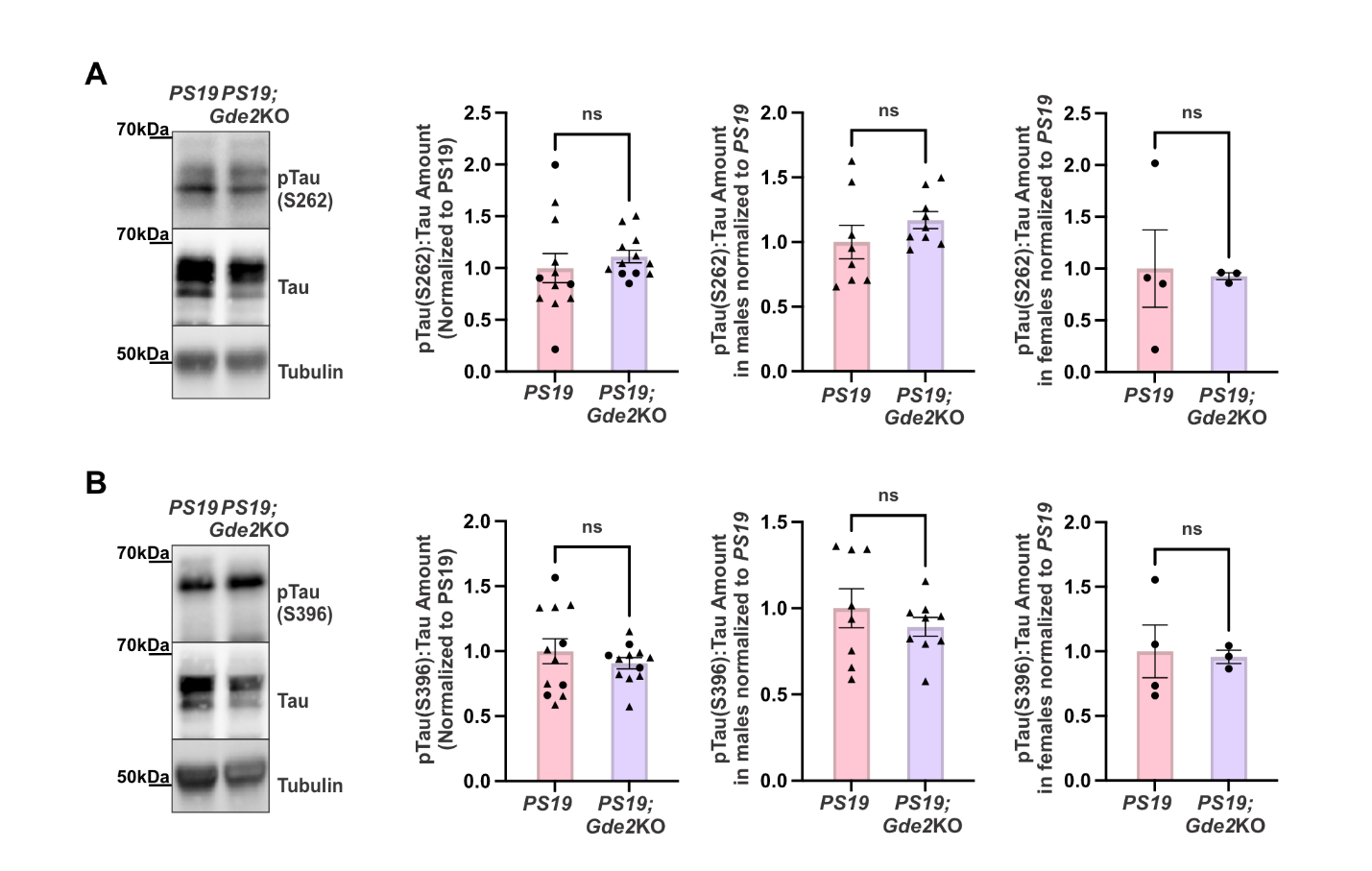
**Fig. S2: Tau phosphorylation at S262 and S396 is unchanged in 6-month *PS19*;*Gde2KO* cortex.** **A,B**, Representative western blots of 6-month *PS19* and *PS19;Gde2KO* cortical extracts and graphs with corresponding quantification for the overall cohort (left), males (middle), and females (right). Triangles and dots denote males and females, respectively. **A**, pTau(S262):Tau ratio; left: ns p=0.4737, middle: ns p=0.2446, right: ns p=0.8740. **B**, pTau(S396):Tau ratio; left: ns p=0.3892, middle: ns p=0.3860, right: ns p=0.8688. All graphs: *PS19* n=12 (8 males, 4 females), *PS19;Gde2*KO n=12 (9 males, 3 females). Mean ± SEM, two-tailed unpaired Student’s *t*-test.

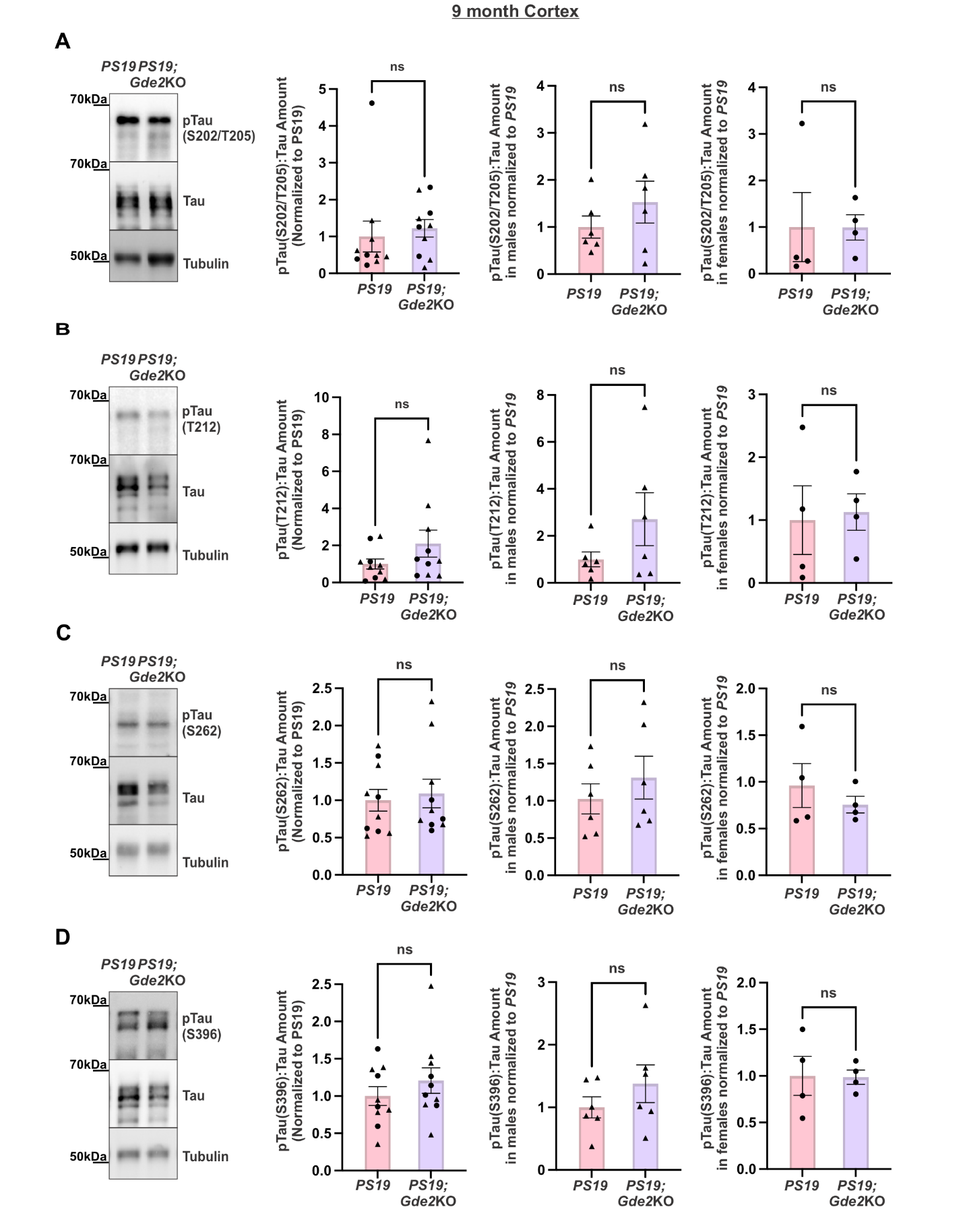
**Fig. S3: Tau phosphorylation is unchanged in the cortex of 9-month *PS19;Gde2*KO animals.** **A-D:** Representative western blots of 9-month *PS19* and *PS19;Gde2KO* cortical extracts and graphs with corresponding quantification for the overall cohort (left), males (middle), and females (right). Triangles and dots denote males and females, respectively. **A**, pTau(S202/T205):Tau ratio; left: ns p=0.6478, middle: ns p=0.3157, right: ns p=0.9939. **B**, pTau(T212):Tau ratio; left: ns p=0.1728, middle: ns p=0.1742, right: ns p=0.8410. **C**, pTau(S262):Tau ratio ns p=0.7132, ns p=0.4346, ns p=0.4456. **D**, pTau(S396):Tau ratio ns p=0.3423, ns p=0.3003, ns p=0.9517. All graphs: *PS19:* 9mo n=10 (6 males, 4 females); *PS19;Gde2KO*: 9mo n=10 (6 males, 4 females). Mean ± SEM, two-tailed unpaired Student’s *t*-test.

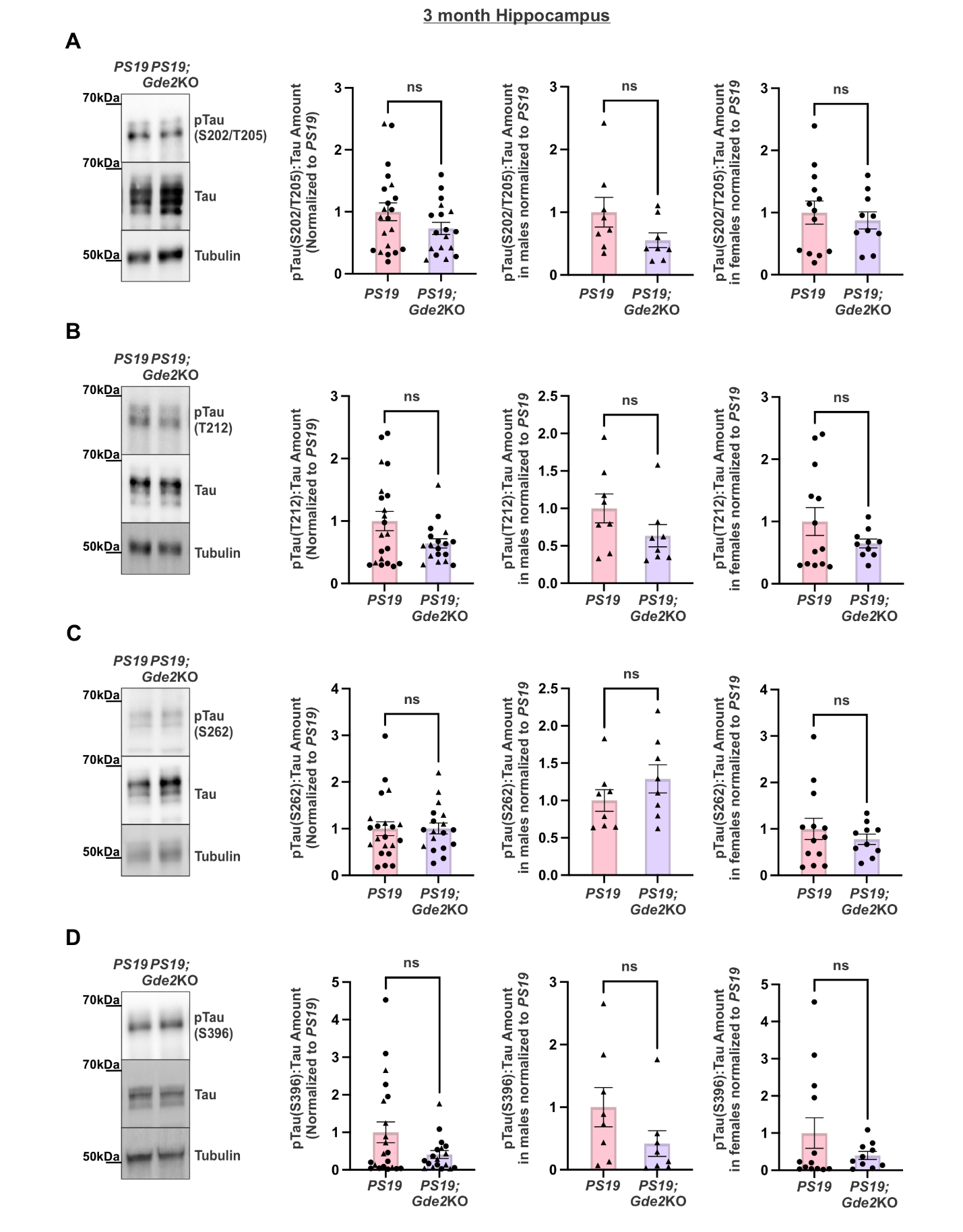
**Fig. S4: Tau phosphorylation is unchanged in the hippocampus of 3-month *PS19;Gde2*KO animals.** **A-D,** Representative western blots of 3-month *PS19* and *PS19;Gde2KO* hippocampal extracts with corresponding quantification for the overall cohort (left), males (middle), and females (right). Triangles and dots denote males and females, respectively. **A**, pTau(S202/T205):Tau ratio; left: ns p=0.1422, middle: ns p=0.1140, right: ns p=0.6116. **B**, pTau(T212):Tau ratio; left: ns p=0.0534, middle: ns p=0.1554, right: ns p=0.1952. **C**, pTau(S262):Tau ratio; left: ns p=0.9800, middle: ns p=0.2442, right: ns p=0.4365. **D**, pTau(S396):Tau ratio; left: ns p=0.0669, middle: ns p=0.1429, right: ns p=0.2252. All graphs: *PS19:* 3mo n=21 (8 males, 13 females); *PS19;Gde2KO*: 3mo n=18 (8 males, 10 females). Mean ± SEM, two-tailed unpaired Student’s *t*-test.

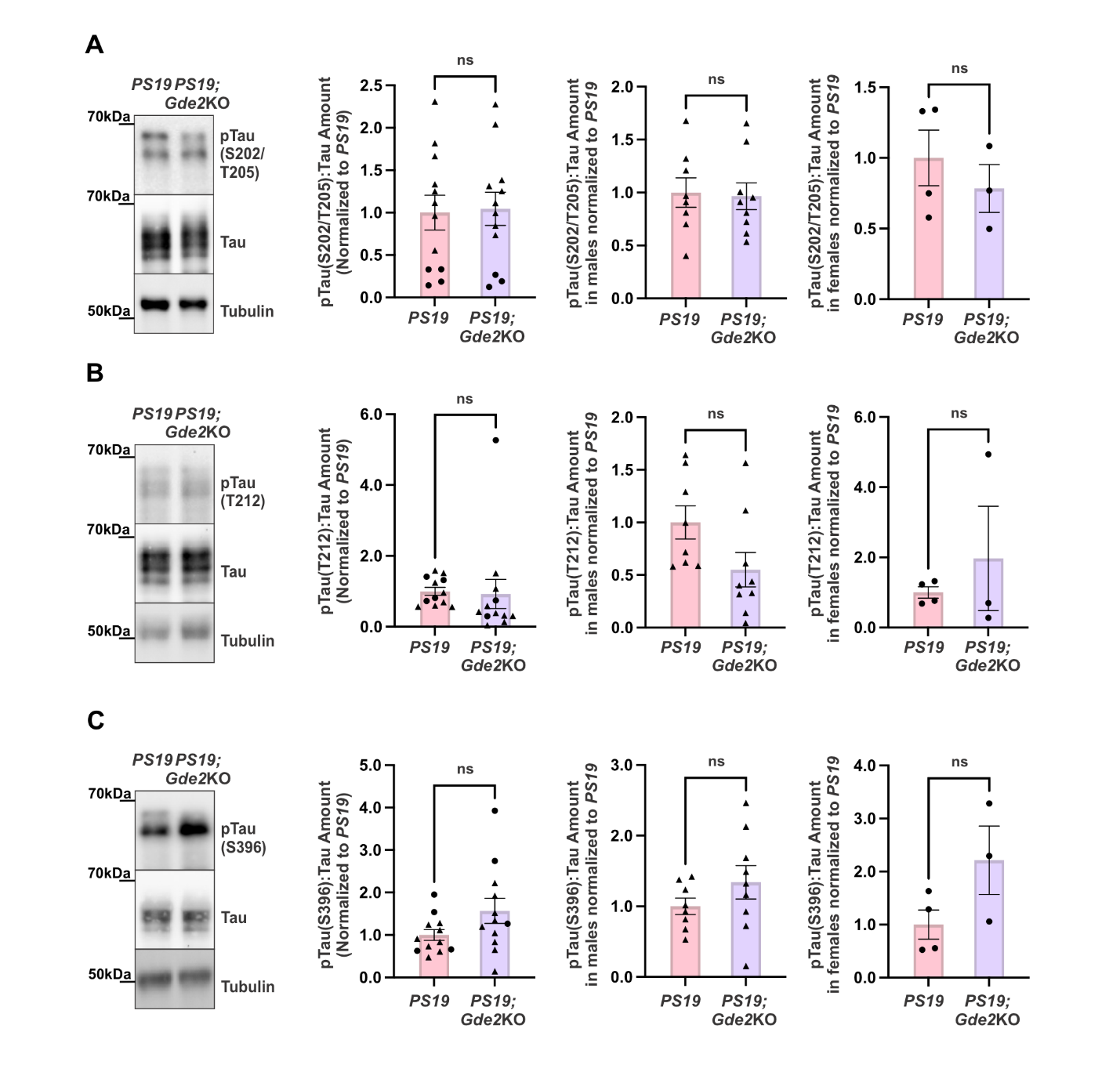

**Fig. S5: Tau phosphorylation at S202/T205, T212, and S396 is unchanged in the 6-month *PS19*;*Gde2KO* hippocampus.** **A-C**, Representative western blots of 6-month *PS19* and *PS19;Gde2KO* hippocampal extracts with corresponding quantification for the overall cohort (left), males (middle), and females (right). Triangles and dots denote males and females, respectively. **A**, pTau(S202/T205):Tau ratio; left: ns p=0.8753, middle: ns p=0.8560, right: ns p=0.4650. **B**, pTau(T212):Tau ratio; left: ns p=0.8622, middle: ns p=0.0677, right: ns p=0.4761. **C**, pTau(S396):Tau ratio; left: ns p=0.0903, middle: ns p=0.2325, right: ns p=0.1119. All graphs: *PS19* n=12 (8 males, 4 females), *PS19;Gde2*KO n=12 (9 males, 3 females). Mean ± SEM, two-tailed unpaired Student’s *t*-test.

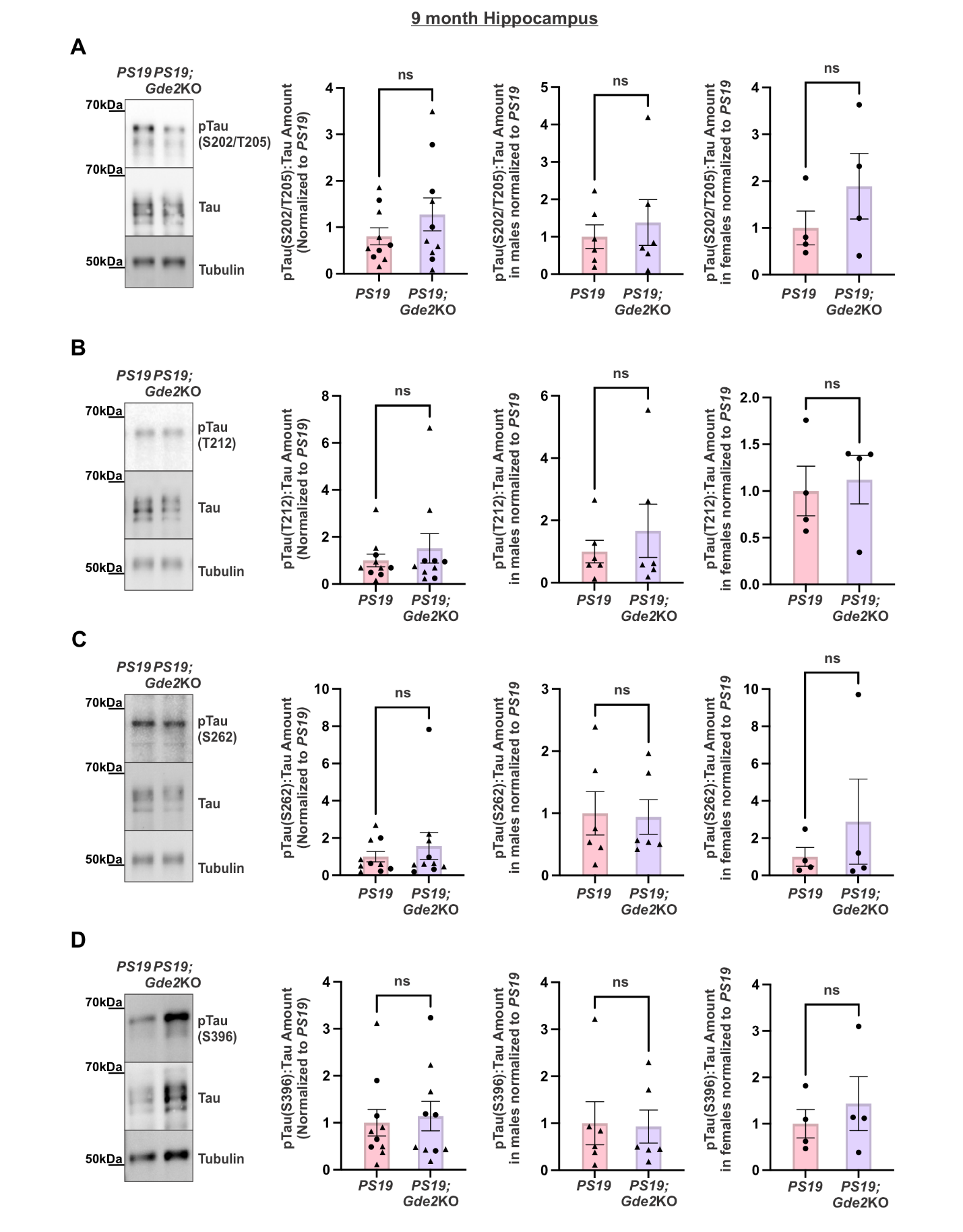
**Fig. S6: Tau phosphorylation is unchanged in the hippocampus of 9-month *PS19;Gde2*KO animals.** **A-D:** Representative western blots of 9-month *PS19* and *PS19;Gde2KO* hippocampal extracts and graphs with corresponding quantification for the overall cohort (left), males (middle), and females (right). Triangles and dots denote males and females, respectively. **A**, pTau(S202/T205):Tau ratio; left: ns p=0.2554, middle: ns p=0.5942, right: ns p=0.3016. **B**, pTau(T212):Tau ratio; left: ns p=0.4621, middle: ns p=0.4892, right: ns p=0.7554. **C**, pTau(S262):Tau ratio; left: ns p=0.4740, middle: ns p=0.8978, right: ns p=0.4507. **D**, pTau(S396):Tau ratio; left: ns p=0.7409, middle: ns p=0.9078, right: ns p=0.5325. All graphs: *PS19:* 9mo n=10 (6 males, 4 females); *PS19;Gde2KO*: 9mo n=10 (6 males, 4 females). Mean ± SEM, two-tailed unpaired Student’s *t*-test.
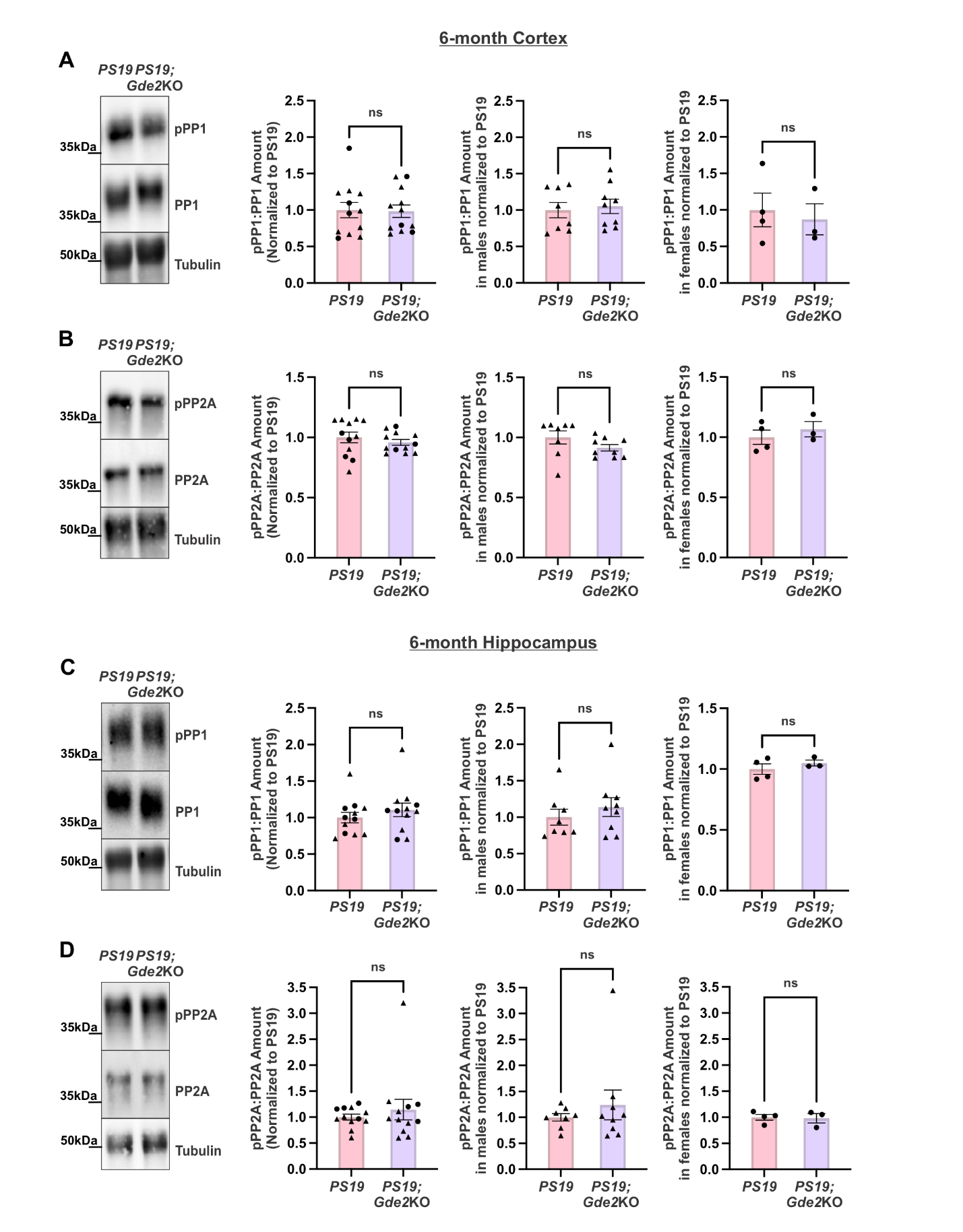
**Fig. S7: Phosphatase activity is unchanged in both the cortex and hippocampus of 6-month *PS19*;*Gde2KO* animals.** **A-D**, Representative western blots of 6-month *PS19* and *PS19;Gde2KO* cortical (**A**, **B**) and hippocampal (**C**, **D**) extracts with corresponding quantification for the overall cohort (left), males (middle), and females (right). **A**, **C**, pPP1:PP1 ratios in 6-month *PS19* and *PS19;Gde2KO* cortical and hippocampal extracts with corresponding quantification. **A**, cortex; left: ns p=0.9102, middle: ns p=0.7184, right: ns p=0.7082; **C**, hippocampus; left: ns p=0.3759, middle: ns p=0.4305, right: ns p=0.4025. **B**, **D**, pPP2A:PP2A ratios in 6-month *PS19* and *PS19;Gde2KO* cortical and hippocampal extracts with corresponding quantification. **B**, cortex; left: ns p=0.4151, middle: ns p=0.1590, right: ns p=0.4851; **D**, hippocampus; left: ns p=0.4908, middle: ns p=0.4551, right: ns p=0.8638. All graphs: *PS19* n=12 (8 males, 4 females), *PS19;Gde2*KO n=12 (9 males, 3 females). Mean ± SEM, two-tailed unpaired Student’s *t*-test.
